# Fluctuations in auxin levels depend upon synchronicity of cell divisions in a one-dimensional model of auxin transport

**DOI:** 10.1101/2023.05.18.541266

**Authors:** Simon Bellows, George Janes, Daniele Avitabile, John R. King, Anthony Bishopp, Etienne Farcot

## Abstract

Auxin is a well-studied plant hormone, the spatial distribution of which remains incompletely understood. Here, we investigate the effects of cell growth and divisions on the dynamics of auxin patterning, using a combination of mathematical modelling and experimental observations. In contrast to most prior work, models are not designed or tuned with the aim to produce a specific auxin pattern. Instead, we use well-established techniques from dynamical systems theory to uncover and classify ranges of auxin patterns as exhaustively as possible, as parameters are varied. Previous work using these techniques has shown how a multitude of stable auxin patterns may coexist, each attainable from a specific ensemble of initial conditions. When a key parameter spans a range of values, these steady patterns form a geometric curve with successive folds, often nicknamed a snaking diagram. As we introduce growth and cell divisions into a one-dimensional model of auxin distribution, we observe new behaviour which can be conveniently explained in terms of this diagram. Cell growth changes the shape of the snaking diagram, corresponding to deformations of auxin patterns. As divisions occur this can lead to abrupt creation or annihilation of auxin peaks. We term this phenomenon ‘snake-jumping’. Under rhythmic cell divisions, we show how this can lead to stable oscillations of auxin. However, we also show that this requires a high level of synchronisation between cell divisions. Using 18 hour time-lapse imaging of the auxin reporter DII:Venus in roots of *Arabidopsis thaliana*, we show auxin fluctuates greatly, both in terms of amplitude and periodicity, consistent with the snake-jumping events observed with non-synchronised cell divisions. Periodic signals downstream the auxin signalling pathway have previously been recorded in plant roots. The present work shows that auxin alone is unlikely to play the role of a pacemaker in this context.

**Author summary:** Auxin is a crucial plant hormone, the function of which underpins almost every known plant development process. The complexity of its transport and signalling mechanisms, alongside the inability to image directly, make mathematical modelling an integral part of research on auxin. One particularly intriguing phenomenon is the experimental observation of oscillations downstream of auxin pathway, which serve as initiator for lateral organ formation. Existing literature, with the aid of modelling, has presented both auxin transport and signalling as potential drivers for these oscillations. In this study, we demonstrate how growth and cell divisions may trigger fluctuations of auxin with significant amplitude, which may lead to regular oscillations in situations where cell divisions are highly synchronised. More physiological conditions including variations in the timing of cell divisions lead to much less temporal regularity in auxin variations. Time-lapse microscope images confirm this lack of regularity of auxin fluctuations in the root apical meristem. Together our findings indicate that auxin changes are unlikely to be strictly periodic in tissues that do not undergo synchronous cell divisions and that other factors may have a robust ability to convert irregular auxin inputs into the periodic outputs underpinning root development.

## Introduction

As a plant hormone playing a key role in virtually every vegetal development process, auxin has attracted a huge amount of research since its discovery at the onset of the 20^*th*^ century [1]. Despite its relatively small and simple structure, auxin affects a very wide range of different responses in plant tissues. Conceptually, this indicates that the complexity of its function lies not in its structure, but in the *processes* it participates in. The prominence of a process over its underlying actors is a viewpoint found at least as early as Heraclitus, and which is still significant in contemporary research [2, 3]. The mathematical theory of processes, known as dynamical systems theory, has been significantly developed over the same period as auxin biology [4]. It is able to describe *qualitative* properties of systems evolving in time, in the sense that they remain true for entire ranges of underlying parameterisations. For example, one may aim to predict whether a system has the ability to oscillate spontaneously, for a range of physiologically plausible conditions, rather than look for specific periods or amplitudes occurring with specific parameter values. This is especially relevant to biology, where parameter values are often not known accurately and/or may vary significantly among individuals or species. As some readers may have limited familiarity with this theory, a brief and informal glossary is provided in **S1 File**.

Auxin is produced and degraded via pathways that are still a topic of investigation [5]. Within cells, auxin triggers gene responses by means of a canonical pathway involving protein-protein interactions, transcription and feedback [6], as well as some quicker non-transcriptional responses [7]. Despite the large number of genes responding to auxin, we will hereafter use the generic term “auxin response” to designate changes in transcription of these genes. Auxin is reallocated over long and short distances by means of a complex transport process involving active transport proteins [8].

Although spatial patterns of auxin have been studied using mathematical modelling, this has largely been restricted to static domains. Yet, important auxin responses take place in growing plant tissues, known as meristems. In addition, auxin patterns are evolving at time-scales which are comparable to growth. Whilst auxin response can be triggered within minutes [9], oscillations of the DR5 reporter, downstream the auxin response pathway, have been observed to oscillate with a period of 6 to 15 hours [10, 11]. Of comparable periodicity, cell divisions within the root apical meristem vary significantly depending on cell sizes and position, within a range of 10 to 53 hours [12, 13].

An important class of models in which growth have been studied are reaction-diffusion models, famously introduced in such contexts by Turing [14]. The effects of growth can be significant [15–21]. However, this is not directly applicable to auxin, as it is actively transported rather than its distribution being controlled purely by diffusion. More precisely, several families of auxin transporter proteins accumulate on sub-domains of cell membranes. Their concentration and location are dependent on the concentration of auxin itself [8]. This process is not fully understood at the molecular level, and a number of hypotheses and mathematical models have been proposed, often classified into the two families of “flux-based” and “gradient-based” (or “concentration-based”) models. As the terms indicate, in the former, transporters accumulate as a function of auxin flux through membranes, whilst the latter sees auxin difference between neighbouring cells as the driving quantity [22]. Historically, flux-based models were believed to be better suited to linear patterns such as veins, whilst gradient-based models were supposedly more natural candidates for spotted patterns such as phyllotactic arrangements [23, 24]. However, more recent studies show that this classification is inadequate, as both classes of model can indeed generate a wide range of patterns, including both spots and stripes [22, 25–27]. The formation of these pattern differs from reaction-diffusion models, where a “flat” steady state becomes unstable and gives rise to patterns as a diffusion coefficient reaches a bifurcation point. In contrast, in a typical flux-based model the homogeneous “flat” steady state is unconditionally stable [28], whereas in gradient-based models it does not always exist [29].

In addition to steady patterns, there is experimental evidence of more complex auxin motion, such as travelling waves or localized oscillations. For instance, there have been repeated observations of oscillatory signals of auxin responsive genes, in a specific *oscillatory zone* within plant roots [11, 30]. There are also experimental records of complex centrifugal waves of high auxin in shoot apical meristems [31]. Such rhythmic processes are related to the self-similar structure of plants, where near-identical organs repeatedly emerge from growing meristems [32]. Using mathematical analysis, oscillations have been shown to occur in both classes of transport models, via a so-called Hopf bifurcation [28, 29], but their shape differs drastically from experimental observation. Travelling waves have also been analysed in [33]. A long-stranding strategy in modelling auxin transport has been to use computer simulations with parameters tuned to produce plausible patterns [34–38], including a signalling module on a growing root template and shows consistency with experimental data [39]. However, that work uses a pre-defined periodic input, rather than showing emergent oscillations. Another group uses computer simulations on a realistic root template to show how patterns of cell divisions are sufficient to induce auxin oscillations, via a mechanism they term reflux-and-growth [40, 41]. Experimental data to date indicates an oscillation in auxin response.

Here we uncover and classify families of auxin patterns, in tissues that grow and cells that divide, using dynamical systems theory. We rely on a well established auxin transport model, where this systematic approach has already been performed in a static context. It is known that auxin transport models can lead to a large number of co-existing patterns. Each pattern corresponds to a different distribution of auxin within a tissue and is attained from a specific ensemble of initial auxin distributions. Tracking these patterns as a parameter is varied leads to a curve, with folds occurring for each possible pattern. The use of the term ‘snaking diagram’ in the literature to refer to these folds. Our **Results** include a review of this snaking phenomenon with a static one-dimensional template. Then, we show how growth and cell divisions induce new behaviour including sudden changes in auxin maxima, which we term snake-jumping as they are explained by overlaying snaking diagrams occurring for different configurations of cells. When cell divisions occur synchronously this can lead to localized oscillations of auxin. By adding some randomness to cell sizes and division events, a loss in the regular periodicity of auxin leads instead to more unpredictable fluctuations. We then present 18 hour time courses of the DII:Venus auxin reporter [42], which are consistent with our numerical results with asynchronised cell divisions. We conclude the paper by discussing the biological implications of our findings.

## Results

### Simulations on a static domain

Before describing the effects of auxin distribution in a growing tissue, we first present the case of a static domain. These simulations will provide a framework for the later sections where we focus on the effects of growth.

The model we use was first published in [34]; it includes auxin concentration, and the concentration and subcellular localization of the PIN transporters on a linear template. The flux of auxin between cells is mediated by both diffusion and active transport. PIN activity is dependent on auxin levels and gradients between cells, similarly to other concentration based models [24, 25, 34, 35]. See **Materials and methods** for equations. Our template comprises around 128 cells (depending on simulations) arranged in single file. One end of the domain allows no flux and represents the root apex, whilst the other end is open ended representing a boundary with the mature root. Essentially this could represent a single file of cells within a root. We have selected this simple template as it allows for more exhaustive analysis including systematically examining the existence of multiple steady states and incorporation of growth dynamics. Our template is much simpler than those used in many previous studies, however it has the advantage that we can more easily modulate cell division. Our research takes a subtly different approach to many previous studies. Instead of attempting to recapitulate experimental observations *in-silico*, we use the model to explore what patterns are possible, and only then look if these can be observed *in-planta*. Note that the key property discussed below, i.e. the existence of a snaking bifurcation diagram, is not specific to this template and persists for instance in two-dimensional domains [29].

We ran simulations using parameter sets listed in **Table 1** but with variations in auxin transport (parameter *T*). To find all steady states as *T* varies, we employed the numerical continuation algorithms from [43], implemented in Matlab in [44]. For low values of *T* (below 1.9), we see a single steady state solution with high auxin throughout the tissue, which we term a proto-peak hereafter (Fig 1**B**). With increasing of *T* (2.1), multiple steady states exist for the same value. In a template of 128 cells, this identified multiple steady states with the numbers of peaks varying from one to nine. These patterns co-exist for a given value of *T* (Fig 1B and **S1 Video**). The addition of a peak corresponds to a separate bifurcation event materializing as a fold in the bifurcation diagram (Fig 1A), with multiple folds termed as snaking. These results are similar to previous observations [29], providing confidence to move to dynamic templates.

**Table 1.** Table of default parameter values. Parameter values are partly arbitrary, chosen both to match previous literature and to avoid physical implausibility. Spatial units are expressed in terms of a typical length scale ℒ, indicative of an average cell dimension, of the order of 10 −100*μm*. Note that the exact value of ℒ does not affect the relative magnitudes of spatially dependent parameters, as seen from their occurrence in the model equations and the table below. In the model without growth, cell volumes are set to 1 (i.e. ℒ).

| Variable | Value | Description |
| --- | --- | --- |
| $\rho_{IAA}$ | $0.85 \text{ (nmol} \cdot \mathcal{L}^{-3} \cdot \text{min}^{-1})$ | Auxin production rate |
| $\kappa_{IAA}$ | $1 \text{ (nmol}^{-1} \cdot \mathcal{L}^3)$ | Michaelis constant for auxin synthesis |
| $\mu_{IAA}$ | $0.1 \text{ (min}^{-1})$ | Auxin decay rate |
| $D$ | $1 \text{ (}\mathcal{L} \cdot \text{min}^{-1})$ | Auxin permeability |
| $T$ | Specified with simulations $(\mathcal{L} \cdot \text{min}^{-1})$ | Active transport coefficient |
| $c_1$ | $1.099 \text{ (nmol}^{-1} \cdot \mathcal{L}^3)$ | Exponentiation base for PIN relocation |
| $\kappa_T$ | 1 (dimensionless) | Auxin transport saturation |
| $\rho_{PIN0}$ | $0 \text{ (nmol} \cdot \mathcal{L}^{-3} \cdot \text{min}^{-1})$ | Base production of PIN |
| $\rho_{PIN}$ | $1 \text{ (min}^{-1})$ | Auxin dependent rate of PIN production |
| $\kappa_{PIN}$ | $1 \text{ (nmol}^{-1} \cdot \mathcal{L}^3)$ | Michaelis constant for PIN production |
| $\mu_{PIN}$ | $0.1 \text{ (min}^{-1})$ | PIN decay rate |
| $g$ | $1/3000 \text{ (}\mathcal{L}^3 \cdot \text{min}^{-1})$ | Growth rate of a cell |
| $V_{max}$ | $4/3 \text{ (}\mathcal{L}^3)$ | Volume at which cells divide or stop growing |
| $A_{ij}$ | $1 \text{ (}\mathcal{L}^2)$ | Surface area inter cells $i$ and $j$ |

**Fig 1.**
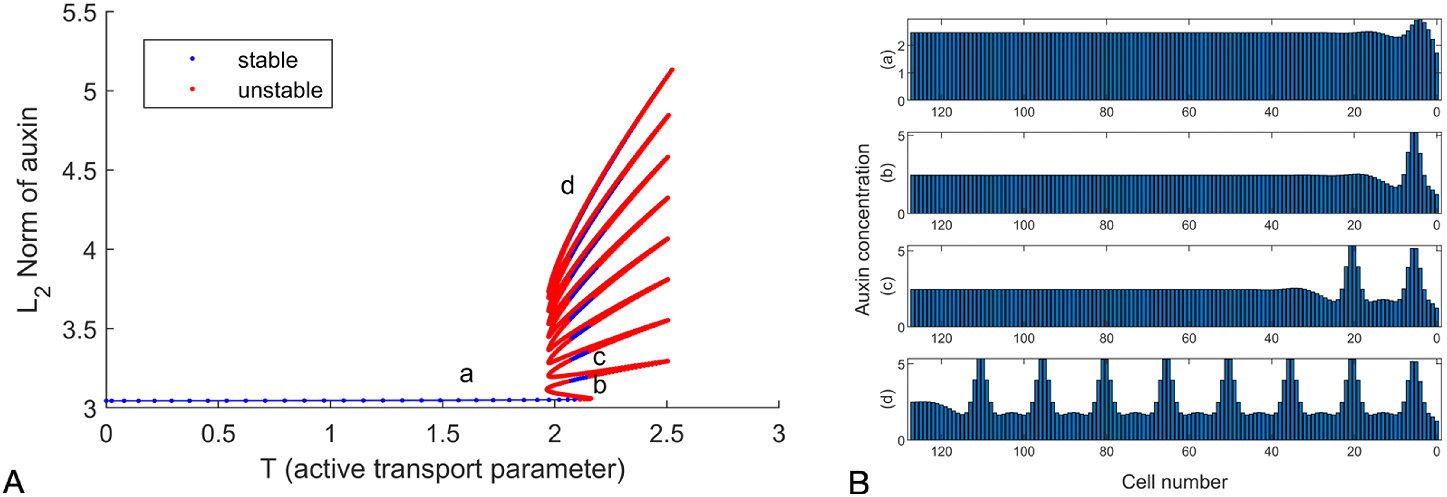
Active transport can lead to multiple stable patterns in auxin concentration. A: The *L*_2_ norm of auxin in the tissue (representative of its total amount) is used in ordinate with active transport parameter T on the *x*-axis. This is sufficient to distinguish multiple solutions (a)-(d), as well as others found as the transport coefficient *T* is used as bifurcation parameter. Shown here is the end result of the analysis, see **S1 Video** for an animated view of the construction of this snaking diagram. B: Example stable solutions marked in A, including the “proto-peak” (a) and 1, 2 and full peaks in a row of 128 cells (corresponding to branches b,c and d on 1A.) for Eqs (1)-(4) with *T* =2.1 and other parameters as shown in **Table 1**.

The steady state solutions can be divided into stable states that are robust to small perturbations in auxin distribution, and unstable states which revert to a stable state upon minor changes in auxin. Although we show unstable solutions in **S1 Fig**, these represent theoretical scenarios: as in any physiological system, even the slightest perturbation (e.g. a 10^−18^ change in auxin concentration) would revert to a stable steady state such as those in Fig 1. These unstable branches are however pertinent to understand the dynamics of the system. They may furthermore gain stability as the domain geometry changes through growth, the topic of next section.

Overall, the number of branches in the snaking region is limited by the length of the domain. In line with this, if the domain were unbounded there would be an infinite number of branches. It should be noted that these branches are themselves shadowed by branches containing solutions of permutations of unstable peaks, as shown in **S1 Fig**. This is likely caused by this taking place on a finite domain, and in [45] this is shown for the well known Swift-Hohenberg reaction diffusion model, specifically when the peaks get within half a wavelength of the boundary. As such, in the present case this systematically occurs, due to the location of the first peak.

### Simulations with deterministic growth

We next tested the effect of growth and cell division upon the snaking diagram. We report results on numerical simulations of the model (5)-(7) on a growing linear chain, using the procedure described in **Materials and methods**. Starting with a pair of cells, we considered a chain in which only the rightmost eight cells could divide. This is a simplification of the Arabidopsis root, in which typically the 30 or so most proximal cells divide. The 8 end cells elongate at constant rate until they reach a specified maximum volume *V*_*max*_, at which point a cell splits into two daughter cells of volume 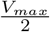. This simple cell division model is well supported by experimental evidence [13]. See **Materials and methods** for complete details. We used the same ODE parameters as previously and reported in **Table 1**, where only the transport rate *T* is left unspecified since we use it as main bifurcation parameter.

Note that for the purpose of visual representation in subsequent figures, cells are depicted as rectangles, with an arbitrary width and a length proportional to the volumes *V*_*i*_ used in the model equations. This does not entail any loss of generality since the model does not depend on a specific cell geometry but only on cell volumes and areas of contact, and only linear chains of cells are considered. Also, as discussed under **Table 1**, units of length can be scaled without effect on the model. Hence the use of any multiple of *V*_*i*_ on the *x* axes of figures is a valid representation of the model geometry.

However, the relative sizes of cells within a domain are expected to have an effect on patterning. Indeed, in the literature on snaking diagrams it has been shown that the precise arrangement of folds and their number depend on the domain geometry [46]. By including growth of the template within our model, one can therefore expect dynamical changes in auxin pattern amongst a wide range of evolving templates.

As cells change in length from their minimal size just after a division event to their maximal size, the snaking diagram shifts and changes in shape (Fig 2A). When cells at the right end of template (representing meristematic cells) are smaller, then the occurrence of a steady state of auxin with one auxin peak is restricted to lower values of *T*. Conversely, as cells elongate the occurrence of the same peak is restricted to higher values of *T*. This confirms that cell size has a profound effect on determining the available auxin maxima within a tissue.

**Fig 2.**
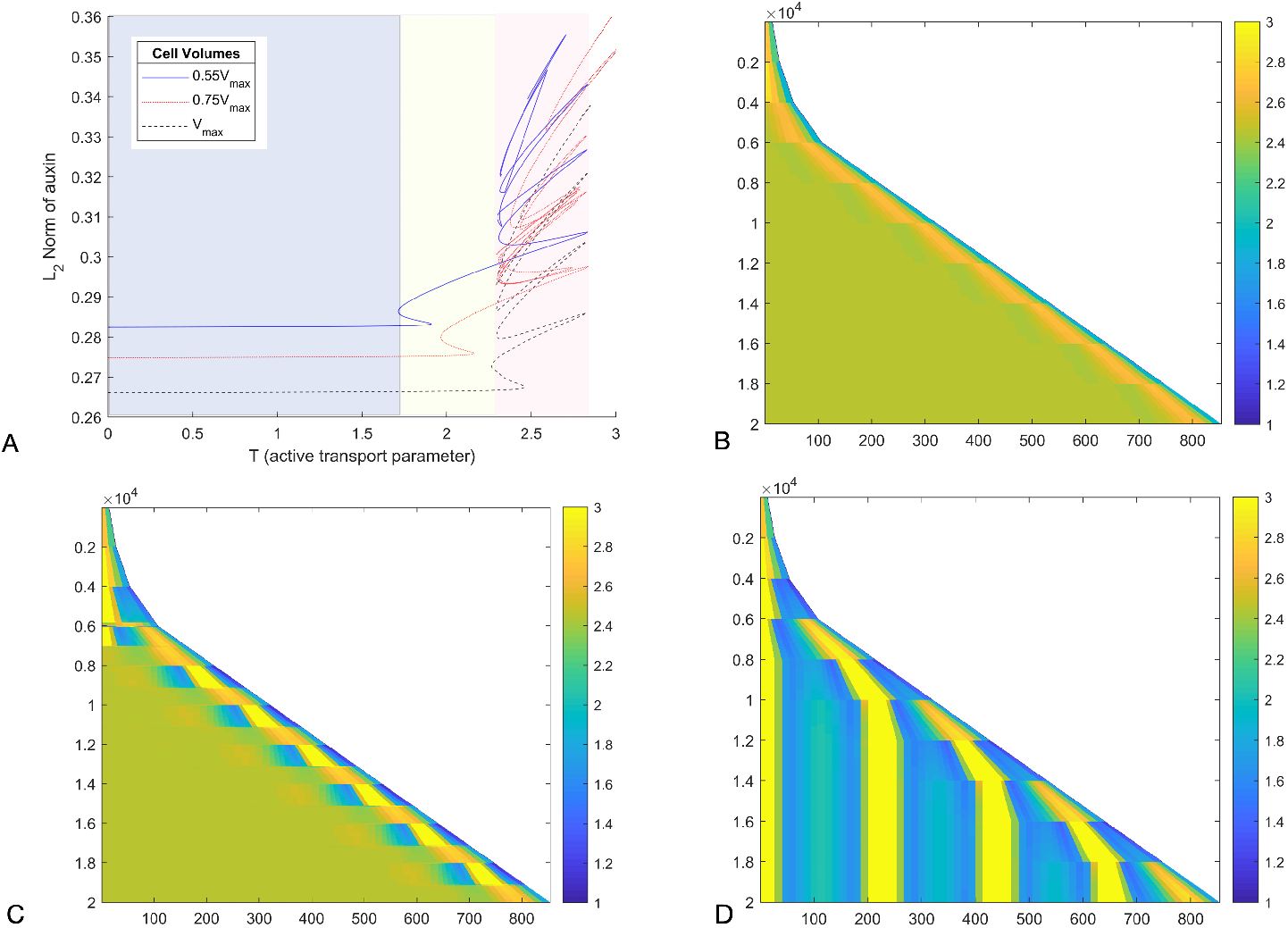
Regular cell divisions produce oscillatory patterns of local auxin accumulation. Growth only occurs for the 8 rightmost cells with the model (5)-(7). A: Continuation in *T*, for three different cell volumes overlaid. Shaded regions correspond to cases 1-3 discussed in the main text. B-D Time-stepping of the same model, with time as *y* coordinate, increasing downwards, for different values of *T*. The *x*-axis represents cell volumes, scaled so *V*_*max*_ =10 (to be indicative of typical cell volumes in pL). B (*T* =1.5, case 1): the proto-peak is maintained despite growth. C (*T* =2.0, case 2): the system alternates between two distinct one-peak solutions. D (*T* =2.5, case 3): new peaks appear over time near the right end of the domain. See also **S2 Video** for animated versions of B-D.

We observed three qualitatively distinct behaviours, as shaded on Fig 2A. These three regimes are reached for increasing values of *T* in the order below:

#### Case 1 (*Low T*)

The system stays locked to a proto-peak similar to the static case shown in Fig 1A(a), see Fig 2B.

#### Case 2 (*Intermediate T*)

The system switches from the proto-peak state to a single peak of auxin, see Fig 2C. The transition from a proto-peak to a single peak can occur at lower levels of *T* than in the static model, due to changes in cell sizes.

#### Case 3 (*High T*)

The system stays locked at a state of maximal peaks with the even expansion of the template causing the creation of regularly spaced new peaks over time, see Fig 2D.

Our categorisation can be systematised by considering bifurcation diagrams. For example, one can see in Fig 2A, that the snaking bifurcation diagram is altered as cell sizes are assigned different values. Therefore, the alteration may correspond to a change in the number of steady states, or simply in their location in state space. At a cell division event, cell sizes are suddenly halved, so that the steady states available in the system correspond to different snaking diagrams out of Fig 2A. Thus, even if the system may not evolve fast enough to be close to a new steady state, the fact that steady states are altered entails a re-direction on the transient dynamics, resulting in noticeable qualitative change. As this is well captured by superimposing snaking diagrams shown in Fig 2A, we casually refer to this general phenomenon as *snake jumping* in the following, as any prior occurrence of the term is from an unambiguously different context [47].

Over time, the cell size *V* varies in the range [*V*_*max*_*/*2, *V*_*max*_]. Therefore, it follows that the bifurcation diagrams (in *T*) obtained for fixed volumes of *V*_*max*_*/*2 and *V*_*max*_ respectively, will bound the set of all snaking diagrams obtained for other values of *V*. As both the *V*_*max*_*/*2 and *V*_*max*_ diagrams have a similar shape to that observed in the static simulations in Fig 1A, it is expected that a single branch will be available regardless of *V* for lower values of *T*. For intermediate values of *T*, this will likely correspond to a coexistence of steady states allowing snake jumping to occur.

The diagrams in Fig 2A illustrate how the first snaking curve occurs for different *T* as cell volume varies. The Case 2 patterns are caused by the state jumping between one and no peaks, as seen in Figure Fig 2C. This is further confirmed by taking *V* as a parameter and continuing solutions: one can see the occurrence of a bifurcation enabling a snake jump from a single full peak to a proto-peak, see **S2 Fig**. Note that for larger values of *T*, the system becomes locked to the fully patterned state, where peaks are spread over the whole domain.

One observation that can be made is that for intermediate values of *T*, near the tip of the domain auxin concentration oscillates. This appears in Fig 2C, and is made more apparent using a video animation of the time-stepping, see **S2 Video**. Oscillations of auxin response in time have been observed in the model plant *Arabidopsis thaliana* [11, 30] *as discussed further in the final section*.

### Effect of irregularity in cell divisions

*A limitation of the simulations discussed in the previous section is that within the dividing zone* 𝒟 *(t*) all cells divide simultaneously, since they are initialised at the same size, grow at the same rate and divide at the same threshold. This is not only unrealistic, but it may invalidate our reasoning about snake jumping inducing auxin oscillations. Indeed, the well separated bifurcation diagrams in Fig 2A correspond to domains where a significant number of cells have a different size. In contrast, one may expect less distinctly separated bifurcation diagrams for uneven distributions of cell sizes with less drastic differences (for instance when cell divisions are asynchronous), obstructing snake jumps.

Thus, given the key role played by cell division, and the fact that the timing of cell divisions and the resulting cell sizes are variable within a growing tissue, we carried out simulations where the size of the daughter cells after a division event included some randomness, see **Materials and methods** for details. In previous analyses we had a scenario in which cells grew linearly and divided when they reach a size *V*_*max*_, at which point they were split into two cells of half that size. Now, the same linear growth and threshold apply, but daughter cells have uneven sizes, chosen randomly. As a result, the distribution of cell sizes and hence the timing division events lose the complete determinism and homogeneity seen in previous simulations.

When we made only modest changes in the synchronicity of cell divisions, we saw patterns broadly similar to the synchronous case above. However, when we introduced a greater level of variability (which we think is more representative of biological templates), we saw irregular and striking fluctuations of auxin.

Hence, the main finding resulting from this study is the variability in cell division is kept low, the occurrence of regular oscillations was unlikely. The correlation between cell size at the onset of division has been studied in the shoot apical meristem, where it has been shown to be highly variable. [48]. In the root meristem, recent studies using live imaging of growing root meristems show that cell divisions do not occur at once, but over a wave of around 10 hours [49]. The rate cell divisions are also variable dependent on cell type and position within the meristem [12, 50]. Though presenting some regularity, all available data on roots indicate a significant variability in the timing of cell division events among neighbouring cells. This is illustrated in our study within Fig 3, where we compare simulation results for two values of the standard deviation of cell ratios after a division event. The low value (*σ* ≈2.3) was the highest value we found to ensure qualitative agreement with the perfect synchrony studied above. This is in contrast with the higher variability (*σ*≈ 9.3), which is based on the only experimental estimates we have found for *σ* in the literature [48]. As seen by comparing Fig 3A and Fig 2C (obtained for very similar values of *T*), the low *σ* simulations are in good (but not complete) agreement with the perfectly synchronised cell divisions. On the other hand, the more realistic dispersion, Fig 3B, leads to a significant irregularity in the changes of auxin peak distribution. This includes short lived travelling waves of auxin, which may appear transiently as an oscillation but have no clearly definable periodicity over the long term (see time 0.6e4 in Fig 3B). As an estimate of periodicity, we computed the distribution of time intervals between successive division events (Fig 3C-D); a positive value, corresponding to emerging periodicity, could be detected for low *σ* only.

**Fig 3.**
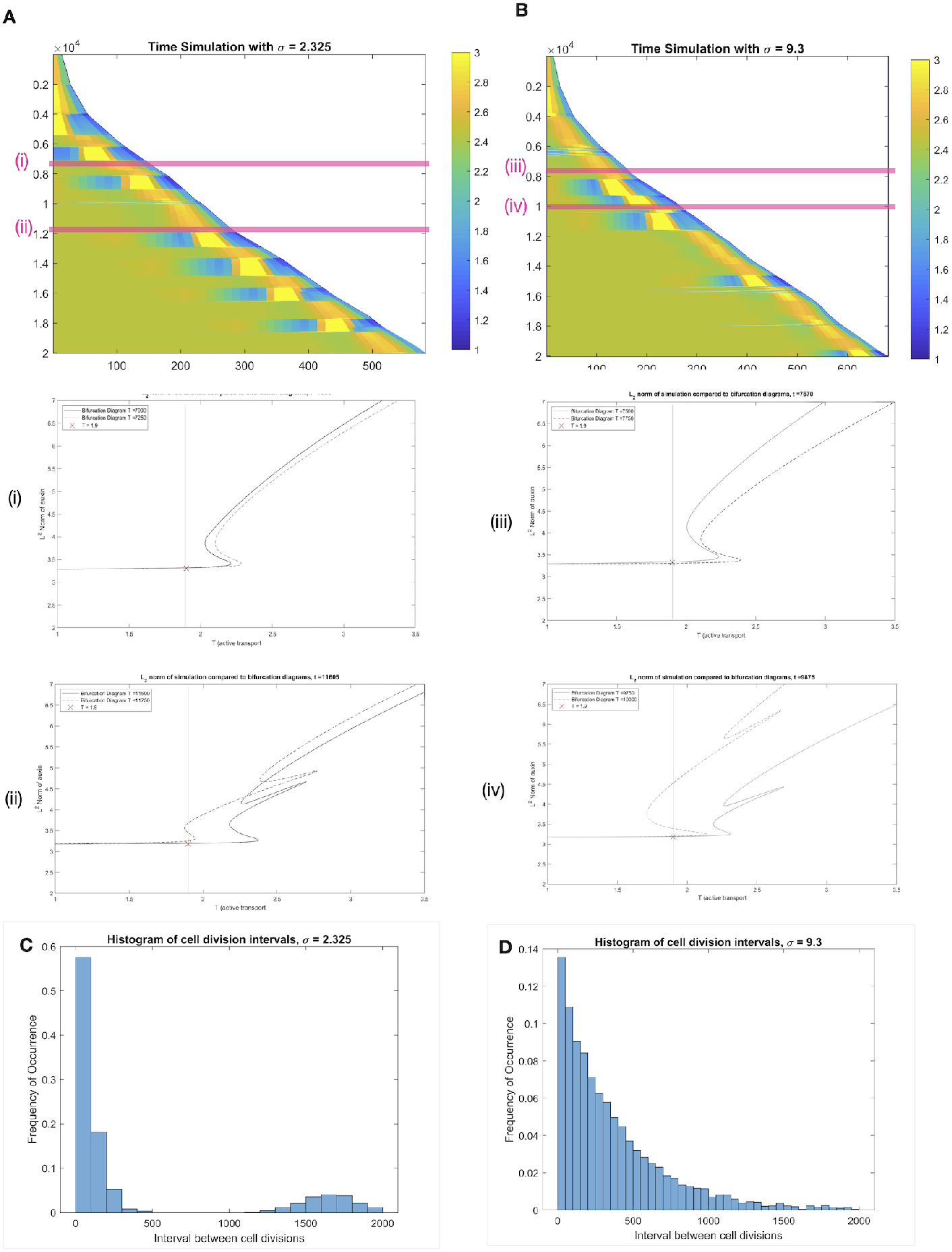
Asynchronous cell divisions trigger irregular auxin fluctuations. Time courses on a growing domain with *T* =1.9, time in *y* and space in *x* coordinates, respectively. Cell sizes after division events are random, with standard deviations *σ* =2.325 (A) and *σ* =9.3 (B). The latter is comparable to experimental data. Histograms (C-D) indicate the distribution of time intervals between division events; a local maximum occurs in C, representative of an emerging periodicity, whereas D has no apparent period. In both cases, transient peaks of auxin coincide with snake jumping events; this is highlighted at two time intervals for each *σ* (pink bands in A and B): no snake jumping occurs over time intervals (i) and (ii), some do over (iii) and (iv). Over the full course of the simulations, the snaking diagrams gets continually distorted by changes in tissue geometry; see **S4 Video** and **S5 Video** for animations, of which panels (i)-(iv) above are snapshots, and **S3 Video** for a deterministic analogue.

Importantly, the underlying mechanism of snake jumping is occurring regardless of *σ*: when a tissue is fixed, for a range of values of the transport parameter *T* there is a snaking diagram, with coexistence of multiple peaked patterns. Any change in the tissue geometry will induce a deformation of this diagram. Hence as the tissue grows and cells divide, the branches available for a snake jumps vary in a less stereotypical and predictable way than for synchronous cell division, where the overall tissue geometry is much more constrained.

### Comparison with experimental data

To explore whether these findings tie with experimental evidence requires temporal data on auxin distribution in a growing root tissue. Our model predicts that in a growing template with asynchronous cell divisions (as would occur naturally), we should not expect to see a regular oscillation of auxin.

As mentioned in introduction, there is solid experimental evidence of an oscillatory zone in plant roots. However these data do not include auxin itself but the DR5 reporter [51], which accounts for the transcriptional response to auxin. This response occurs downstream a signalling pathway involving a complex series of nonlinear processes and feedback loops, and has been previoulsy described using differential equation models, see e.g. [52, 53]. Both this modelling work and experimental data indicate that the outputs of the pathway may differ significantly from its auxin input.

Since all results presented above concern the distribution of auxin, without any representation of the signalling pathway, it was difficult to make a direct comparison with DR5 data. The DII:Venus [42] fluorescent reporter is directly degraded by auxin. As such it is more directly representative of auxin concentration than DR5, and therefore a better candidate to support or discount the claims of the previous sections.

On the other hand, while data in e.g. [11, 39] relied on the DR5:LUC construct, which allows live time-lapse imaging over long periods, DII:Venus requires a confocal microscope. This provides higher spatial resolution in which auxin (using DII as a proxy) can be quantified in individual cells. We imaged DII expression in 8 *Arabidopsis* roots at 1h time intervals for 18h, and quantified the fluorescence, see **Materials and methods**. These experimental time series are reported on Fig 4.

**Fig 4.**
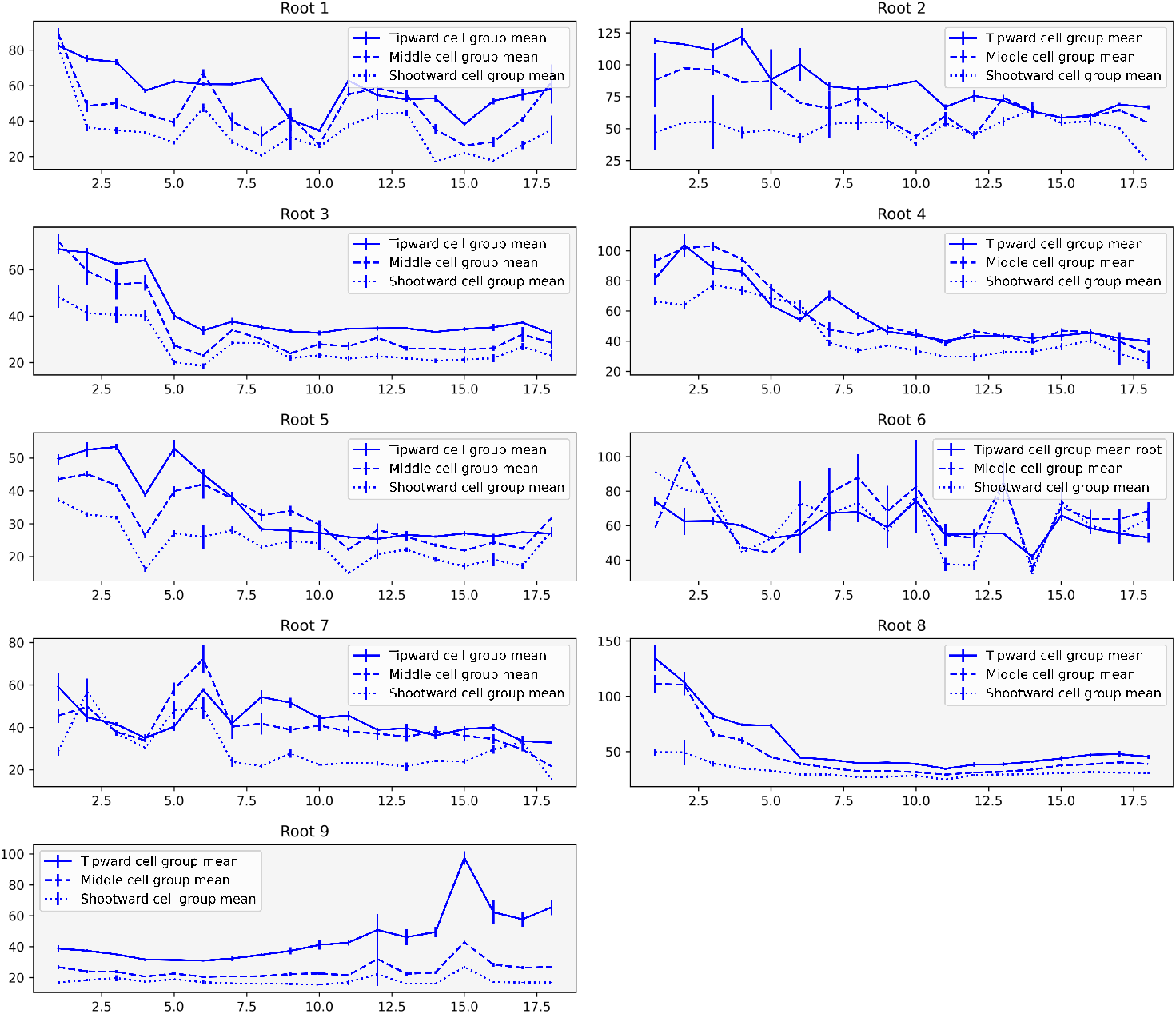
Time series of DII:Venus data. On each graph the three curves represent three successive groups of five cortex cells from the root tip. Curves represent the average of the five cell groups (with three replicates, see text), and error bars the corresponding standard deviation. See protocol in **Materials and methods** and full data set in **S3 File**

We did not observe a clear periodic signal in any of the roots that were tracked. We note almost every root there undergoes noticeable fluctuations over time and space. The temporal profiles show sharp periods of increase and decrease of the fluorescent signal, which can be localized within the tissue. These fluctuations are reminiscent of the simulations performed in the previous section and therefore we attempted a more careful comparison.

The temporal profiles show sharp periods of increase and decrease of the fluorescent signal, which can be localized within the tissue. These fluctuations are reminiscent of the simulations performed in the previous section and therefore we attempted a more careful comparison. In Fig 5, we report some selected simulations as well as DII:Venus time series, see. Note that, due to DII:Venus being an inverse reporter degraded by auxin, the time series were ‘reversed’ (i.e. taking the difference between the maximum value over time and the full time course, for each time course). This matching was obtained by trial and error, relying on the randomness in the simulations: a long time-stepping simulation was compared by eye to the DII:Venus data. Although a more robust statistical comparison would have been preferable, the comparable time scales between auxin fluctuations and the longest feasible experimental set-up forced us to resort to this more qualitative approach. In fact, most time series we produced proved to present the simulated and experimental data sets.

**Fig 5.**
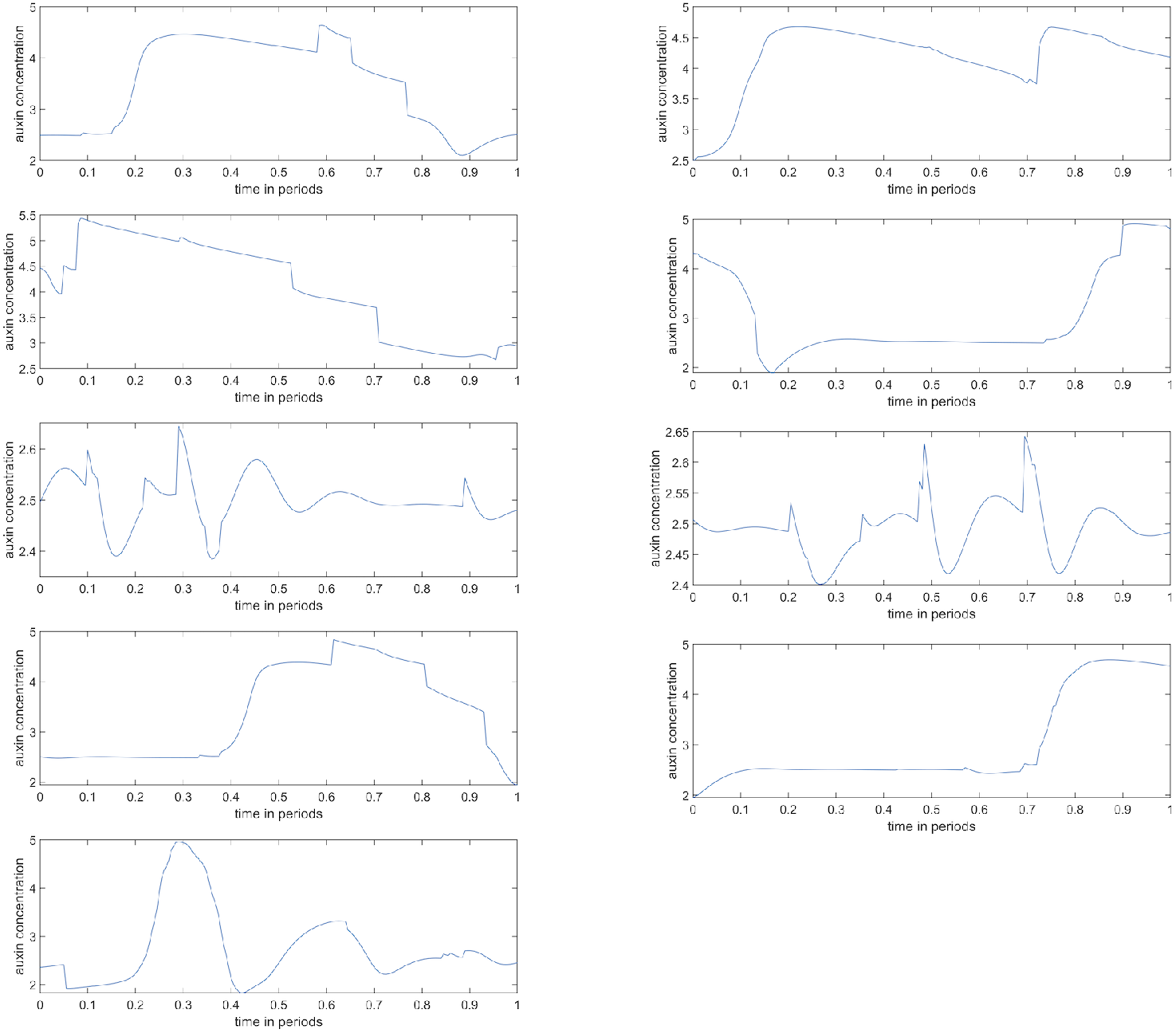
Simulations in qualitative agreement with DII:Venus data. Horizontally: time, normalised to a ‘period’ of 200 time units. Vertically: mean auxin concentration for cells 6 to 10 over the reference time period. See full time series data and further details in **S3 Fig**. The parameters for the simulation are as in Table 1, with the exception of cell growth rate being increased by a factor of 10 to allow for longer time range, cell size variability *σ* =9.3 and *T* = 1.9.

## Discussion

All the results presented above aim to clarify behaviour of an established auxin transport model on growing domains. A key technique in this endeavour was numerical continuation (see **S1 File** for background and **S2 File** for our implementation), whereby a user systematically scans a full range of possible auxin patterns rather than running individual simulations. The active transport coefficient *T* is used as bifurcation parameter on “snapshots” of the domain at specific time points during growth. This shows a specific set of stable steady states (auxin patterns) for each time point, forming a snake-like curve as *T* varies. Each fold of the snaking structure corresponds to a pattern and all patterns compete, attracting distinct sets of initial conditions. Over time, changes in this snaking curve induce changes in which particular pattern attracts the dynamics, leading to potential alternations of patterns across time. We chose to term these alternations snake jumps in reference to the underlying bifurcation diagrams.

In cases where cell divisions are highly synchronised, the number of tissue configurations is strongly limited: all possible tissues are essentially related to each other by a scalar factor (the size of growing cells), ranging over a compact interval. In this scenario, the two extreme values for this factor (just before and after division) provide only two snaking diagrams. Yet, the high number of branches and their associated auxin patterns lead to a range of possible behaviours. In a typical scenario, we observed an alternation between a single auxin peak and none. Interestingly, there have been observations of controlled cell proliferation in roots, which may increase the synchrony of cell divisions in specific tissues [54]. Similarly, the recently proposed mechanism of “reflux-and-growth” [40, 41] shows how computational simulations on realistic tissue geometries can produce an oscillatory zone under the control of tissue growth.

However, in our simulations, the level of synchrony required to induce robust oscillations proved to be typically much higher than the experimental figures we have found in the literature. By running simulations with plausible variations in cell division sizes (and hence timings), we found again that snake jumping occurs and can lead to alternations between qualitatively distinct auxin maxima. However, the number of distinct snaking diagrams now becomes driven by a stochastic process that leads to an infinite number of configurations, albeit qualitatively similar. This additional complexity makes it impossible to decipher any regular pattern in auxin fluctuations. Yet, fluctuations of significant amplitude did typically occur.

This prompted us to produce some experimental validation, which we obtained by means of the DII:Venus reporter. In our experimental time courses, we did indeed observe the same type of irregular auxin fluctuations. We were able to find some surprisingly good qualitative agreement between simulations and experiment. As with any modelling approach, it is inherently impossible to confirm that our modelling proposal is ‘true’. But we can state an excellent consistency between the model predictions and the data. This is largely made possible by the qualitative nature of our predictions, which stems from the use of bifurcation methods.

From the present work and others, it appears that the combination of active transport and cell division can lead to a wide range of auxin signals at the single cell level; typically one expects individual cells to perceive neither a steady level of auxin, nor a very regular oscillatory signal. This stands both for modelling and experimental observations. Since ultimately the role of auxin in plants is to trigger responses in individual cells, this general observation naturally leads to the question of how would such complex signals be transduced by the auxin pathway. This is especially intriguing when considering the fact that this pathway is able to oscillate spontaneously with constant auxin inputs, at least in theory: a Hopf bifurcation was found in [52] to give rise to stable oscillations in a differential equation model. In fact, it has been suggested that the DR5 oscillations occur downstream of auxin itself: in [39] the authors include oscillations as an input to an ODE model rather than having them emerge spontaneously, and provide numerical evidence that periodic auxin waves can enhance these oscillations, if they are in phase. The mechanism uncovered here shows how periodic auxin fluctuations can result from the patterns of growth of the tissue. This opens up the possibility for a complex interplay between rhythms of auxin itself and the protein network it controls. Ultimately, fluctuations of auxin are dependent on the synchronisation between cells undergoing division. In other words, it is the timing of cell mitotic cycles and the spatial distribution of their phase that determine fluctuations of auxin. The source of regular oscillations can be sought at the level of this phase distribution, as well as downstream of auxin, via its signalling pathway.

## Materials and methods

### Auxin transport model

#### Main differential equations

As mentioned earlier, though the notations are slightly adapted, the model presented in this section is based on previous research first published in [34], in particular relying on the bifurcation study performed for this model in [29].

We consider the active transport of auxin in a tissue composed of *N* = *N* (*t*) cells at time *t*. Each cell *i* ∈ 𝒩 (*t*) ≐ {1 … *N* (*t*)} has neighbours 𝒩_*i*_ = 𝒩_*i*_(*t*) ⊂ 𝒩 (*t*), therefore inducing a graph *G* = *G*(*t*) with node set 𝒩 (*t*) and edges (*i, j*) whenever cells *i* and *j* are in physical contact, i.e. *i* ∈ 𝒩_*j*_ or equivalently *j* ∈ 𝒩_*i*_. We denote by *V*_*i*_(*t*) the volume of cell *i*, and *A*_*ij*_(*t*) the exchange surface area between cell *i* and *j*. Please note that whenever it is not the main point of the discussion, one shall drop the dependence on time to simplify notations.

To describe the dynamics of auxin transport, we use *a*_*i*_ to denote the concentration of auxin and *p*_*i*_ to denote the concentration of transporter proteins (PIN), in a cell *i*. Thus, any variation of auxin *a*_*i*_ in cell *i* is due to the interplay between four processes:

- auxin synthesis, whose metabolism is simplified into a Michaelis-Menten rate with saturation constant *ρ*_IAA_ and Michaelis (i.e. half-maximum) constant *κ*_IAA_.
- auxin degradation, with a fixed rate *μ*_IAA_,
- free diffusion towards neighbouring cells *j* ∈ 𝒩_*i*_, described using Fick’s law: *D*(*a*_*i*_ −*a*_*j*_), where *D* is a permeability coefficient, indicative of diffusion between neighbouring cells,
- active transport by the transporter (PIN) proteins *T* (*P*_*ij*_*h*(*a*_*i*_, *a*_*j*_) − *P*_*ji*_*h*(*a*_*j*_, *a*_*i*_)), where *T* is a transport efficiency coefficient, *P*_*ij*_ is the concentration of PINs in cell *i* near the interface with cell *j* and *h*(*x, y*) a function taking different forms in the literature (cf. e.g. [22, 23]), see below for the version from [34].

For each cell *i* ∈ {1, …, *N*} in the fixed geometry model

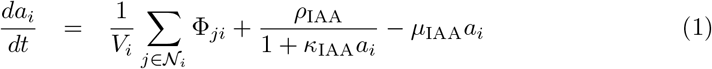

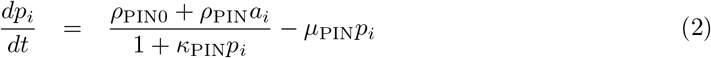

where Φ_*ij*_ denotes the flux of auxin from cell *i* to cell *j*:

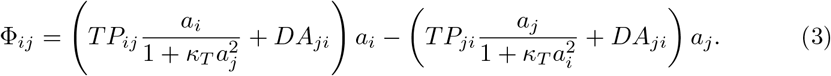

and

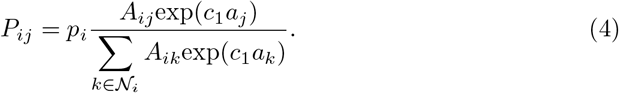

Throughout the paper, the domain consists of a single file of cells with homogeneous contact surface (normalized to 1), and the following neighbouring structure

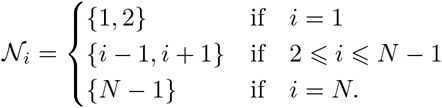

This choice ensures that cell 1, being its own neighbour, mimics the shootwards end of a root as an open-ended interface, where Φ_11_ = 0 from (3), but with a PIN polarity consistent with there being further cells to the left. On the other hand, the only neighbour of *N* is *N* −1 and there is neither flux nor PINs to the right, as would be expected at the root tip.

#### Implementing growth

Whilst understanding how patterning occurs on a fixed domain is important, many realistic dynamical systems occur on growing domains. In biological tissues, growth is typically comprised of cell elongation and cell division. Specifically in plant tissues, these can occur over timescales comparable to auxin variations. Indeed in plant roots cell divisions occur with a typical period of 10-53h hours in Arabidopsis [12, 55].

Periodic oscillations of auxin response are also observed to have a period of a few hours, e.g. 15 hours [30] or 6 hours [11], and are critical to the formation of lateral roots. Hence, considering a growing tissue is relevant not only for mathematical purposes, but is also required by the underlying biology. Furthermore, given the nature of auxin transport, with asymmetric membrane-bound transporters, continuum approximations are unpractical and cell divisions are best represented explicitly.

Regardless of the exact growth model, we must alter the ODEs to take into account dilution due to volume change. Note that concentration is quantity over volume, therefore 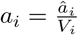, where *â*_*i*_ is the quantity of auxin in cell *i*. Differentiating *a*_*i*_ with respect to time, *t* gives us:

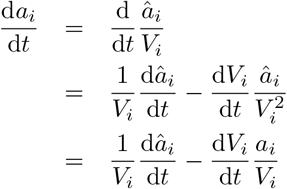

Therefore, a term 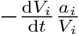 must be added to the right-hand side of Eq (1). Similarly, 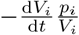 must be introduced in (2).

To mimic growth patterns in roots, one restricts growth to a sub-domain near the tip of the tissue [55]. Note that for theoretical purposes, growth over the whole domain has been considered in [56] and, though not reported in detail here, leads to similar mathematical conclusions as shown here.

We denote the set of dividing cells by 𝒟 (*t*) ⊂ 𝒩 (*t*), and more precisely 𝒟 (*t*) = {*N* (*t*) −*N*_*div*_ … *N* (*t*)}, for a fixed *N*_*div*_ representing the size of the growing zone. To retain simplicity we considered growth to take place linearly over time, i.e. at a constant rate denoted by *g*. As in previous simulation studies [34, 37], cells divide when their volume reaches a threshold, here denoted *V*_*max*_. This is in reasonable agreement with experimental data [13]. Thus, the resulting extension of Eqs (1)-(2) with growth and divisions is:

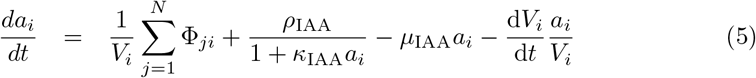

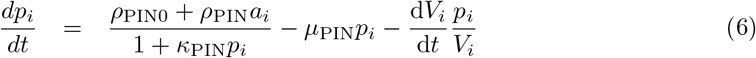

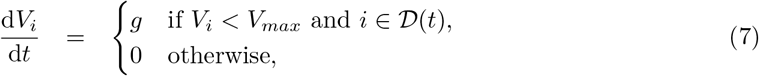

with (3)-(4) still used for the fluxes Φ_*ij*_. Note that the actual cell subscripts, and those of their neighbours, have to be updated after each division event. Therefore, we would need to specify an algorithm for an arbitrary tissue here. For a linear chain of cells, it is natural that, at any time, cell *i* is the left neighbour of cell *i* + 1. When a cell divides, its two “daughter cells” are by default initialised with a volume 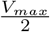, where auxin and PIN concentrations are equal to that of the parent cell and all cells to the right of a division have their subscript increased by one.

Throughout the paper, the parameter values from Table 1 are used.

#### Implementing noise on cell divisions

As real life cell divisions are not expected to occur synchronously, we also implemented a stochastic variant of the previous section, where instead of being 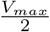 the size of the daughter cells were normally distributed between 24% and 76% of their mother’s size. More specifically, the percentage for one of the daughter cells was set to 50+*x*% where *x* follows a centred normal distribution with variance *σ*^2^, truncated to enforced |*x*| *<* 26 (using the `truncate` routine in Matlab). We considered different values for *σ*, with the highest being derived from a study of cell division in the shoot apical meristem [48] in which a histogram of the proportion daughter cells were of their parents size was a bell curve with a reported standard deviation of *σ* = 9.3.

### DII:Venus confocal imaging

We acquired 18 hourly observations, for nine roots of *Arabidopsis thaliana* grown in identical conditions, see [57] for more experimental background.

#### Plant growth and preparation

DII:Venus seeds cloned using the DEAL vector were sown on 0.5x Murishage and Skoog basal salts with 1% MES monohydrate, 1% agar, pH 5.7. Seeds were then stratified at 4°C in darkness for 2 days before being placed in a growth chamber at 21°C with a 16h light, 8h dark regime. 3-4 days after germination and 12 hours prior to imaging, seedlings were transferred to glass-bottomed imaging cuvettes and the root covered with a slice of growth medium gel. They were then returned to their growth conditions overnight.

#### Imaging and image analysis

Being kept in growth chambers in between imaging sessions, each seedling was imaged once per hour for 18 hours using a Leica SP8 laser scanning confocal microscope with a 40x dry objective. Excitation of both VENUS and tdTomato was performed using a 514 nm laser, and emission collected with photomultiplier tube or hybrid detectors across 520-550 nm and 570-760 nm, respectively. Measurements of fluorescence intensity were made using FIJI software to isolate individual nuclei and output mean gray values.

Three of the authors then recorded manually, for each root, three groups of five cells from the cortex, from the tip upwards, for each time point. This was performed using standard selection and measurement tools in Fiji [58]. We aggregated the three data sets to account for variations between the three records, which were unavoidable due to the experimental noise present in some images, making the boundary of a cell or nucleus partly blurry.

## Supporting information

**S1 Fig. Bifurcation diagram: unstable branches**. A refined and zoomed version of Fig 1 showing additional, unstable branches.

**S2 Fig. Bifurcation diagram with cell volume as control parameter**. As a complement to Fig 2, continuation using cell volume as control parameter can be used to illustrate the creation/annihilation of auxin peaks leading to an oscillatory signal. Axes labels are self-explanatory, *T* = 2.

**S3 Fig. Full time series for Fig 5**, as a heatmap; same legend as Fig 2 and Fig 3. The plots from Fig 5 were obtained by extracting intervals of 200 time units (min) from this time series with random cell division (with *σ* =9.3). The graphs were generated by randomly picking start points throughout the simulation such that they would not overlap with any other interval and plotting the mean auxin over cells 6-10, for the selected 200min interval.

**S1 File. Dynamical systems glossary**. Some background on dynamical systems theory, presented for non-mathematicians.

**S2 File. Matlab code**. The code used for all time-stepping and continuation methods; the provided archive also requires the publicly available Matlab files from [44], which come with a general documentation.

**S3 File. Processed imaging data**. The measured fluorescence data as an excel file. The first three columns indicate the root, domain (out of three groups of 5 cells) and time point recorded. The remaining columns record the mean DII intensity, its standard deviation (within the domain) and the surface area of the selected domain, with three repeats for the three authors replicating the analysis.

**S4 File. Raw images**. Confocal images are available using the following permanent link: https://doi.org/10.17632/p7ftp5wm3h.1.

**S1 Video. Stable patterns along the snaking diagram**. The stable patterns of auxin, including those in Fig1 1A, are displayed as the animation follows the steps of the continuation algorithm underlying the making of Fig 1B.

**S2 Video. Auxin oscillations induced by cell divisions**. Animation representing the simulated growing root with synchronized cell divisions from Fig 2B-D; all three values of *T* are superimposed, with legend within the animation.

**S3 Video. Deformations of snaking under tissue growth: deterministic case**. Animation representing the deformations of the snaking diagram as the domain grows. The crosses follows the time course simulations from Fig 2B-D, with corresponding *T* values in legend.

**S4 Video. Deformations of snaking under tissue growth: low** *σ*. Animation representing the deformations of the snaking diagram as the domain grows as in Fig 3A; *T* =1.9, *σ* =2.325. The red cross follows the time course simulation, which departs transiently from equilibrium branches when they are rapidly updated.

**S5 Video. Deformations of snaking under tissue growth: physiological** *σ*. Animation representing the deformations of the snaking diagram as the domain grows as in Fig 3A; *T* =1.9, *σ* =9.3. The red cross follows the time course simulation, which departs transiently from equilibrium branches when they are rapidly updated.

