## Supplementary figures and images for "Fluctuations in auxin levels depend upon synchronicity of cell divisions in a one-dimensional model of auxin transport"

### S1_Fig

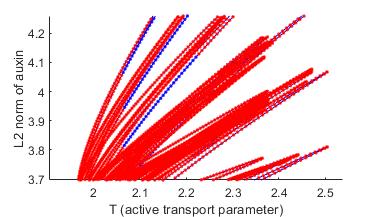

### S2_Fig

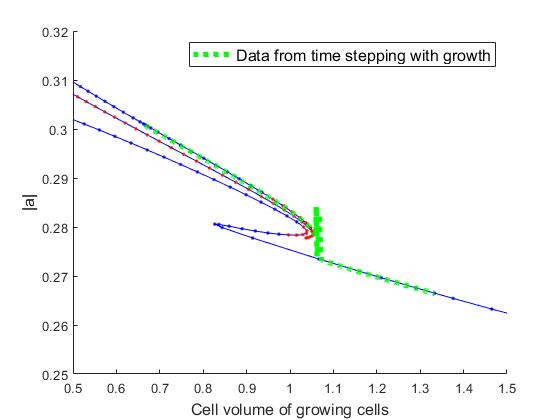

### S3_Fig

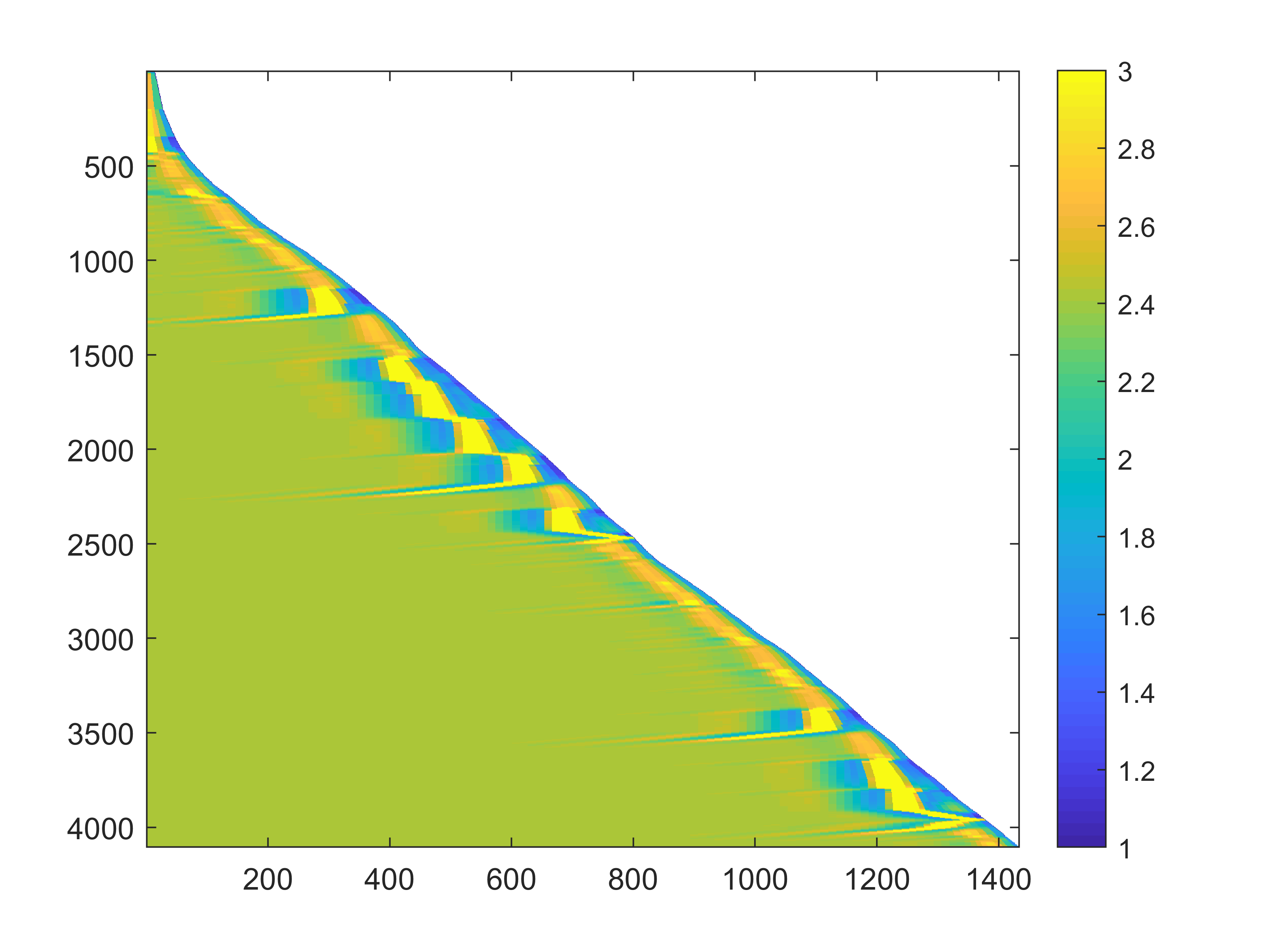
