## Supplementary material for "Fluctuations in auxin levels depend upon synchronicity of cell divisions in a one-dimensional model of auxin transport": S1_File

### S1 File: Dynamical systems: an elementary glossary.

In this section, one avoids technical notations as much as possible and use vernacular language mostly. Terms which have a formal definition in mathematics are highlighted, but all definitions are given using intuition rather than formalism. Readers interested to study more are invited to consult, e.g. [1, 2, 3].
Mathematically, a **dynamical system** consists of two items: a **state space**, describing all possible states (or configurations) for the system of interest, and a **flow**, which is a transformation of that space depending on time. In any example relevant to this paper, time is represented using real numbers, which we denote by $t \in \mathbb{R}$, and the state of the system is represented by a list of nonnegative real numbers, known as **variables** (typically concentrations), which we can denote by $x = (x_1, \ldots, x_n) \in \mathbb{R}^n_{\geqslant 0}$ when there are $n$ variables. In the present context (and in most modelling papers) the flow is not described explicitly; instead, one specifies a system of $n$ differential equations, which encode the inter-dependencies between the $n$ variables $x_i$. The flow is then the transformation which, starting from any initial state $x^0 = (x^0_1, \ldots, x^0_n)$ computes later states $x^t$, for any time $t$, using the differential equations with initial condition $x^0$. In practice this cannot be done using explicit formulas, but can be estimated numerically using well established **time-stepping** methods.
One of the key principles underpinning dynamical systems theory is that the behaviour of a system, and especially its **asymptotic** behaviour (i.e. what happens after a very long time) can be described in geometrical terms. Indeed, as $t$ varies the flow describes **trajectories** which are exactly curves located in state space; when using differential equations they are often termed **solution curves**. The dynamics can then be interpreted in terms of the organization of a system of curves. This is achievable because although there are infinitely many curves, asymptotically all tend to accumulate around a finite number of special regions in state space called **attractors**. In other words entire groups of initial conditions end up following a similar path in state space. Attractors can usefully be classified into a small number of types.
The simplest attractors arise when a single point remains unchanged over time, i.e. the flow remains at that **equilibrium point**, also known as **steady state**, for all times. In such a case, the solutions curve is reduced to a single point. This corresponds to situations that can sustain themselves exactly as they are for indefinitely long periods of time. Importantly, a steady state is physically meaningful only if it is **stable**. In contrast, **unstable** steady states can occur (in which case they are not attractors). For instance, when two genes coding for transcription factors repress each other, one often see three steady states: two with one gene highly transcribed and the other gene "off", and a third where both genes are transcribed at an intermediary level. Only the first two are stable; indeed the intermediary state can not sustain any change, however small, in either gene's expression levels. In the context of this paper, steady states are typically spatial patterns of auxin. More general attractors include periodic solutions, which repeat themselves over time, and form closed curves in state space called **limit cycles**. More complex attractors also exist but are beyond the scope of this paper.

A very important notion in this framework is that of **bifurcation**; besides state variables, dynamical systems typically include **parameters**, which are quantities remaining fixed over time. Examples include reaction rate constants, diffusion rates, half-life constants etc. Though fixed for a given system, parameters may vary between individuals or due to changes in experimental conditions. Mathematically, one can track how the organization of state space gets modified as a parameter is varied. Abrupt changes of dynamics may occur at specific, critical values of the parameter; typically, an attractor appears, disappears, or changes in nature. Such changes are termed bifurcations. Bifurcation theory investigates these changes, again using geometric methods. Looking at a steady state (for example) and varying a parameter value, one builds a curve, known as **branch** of steady state values. Bifurcations correspond exactly to situations where branches of solutions present a fold (hence disappear for some parameter values), or collide and lead to changes in the number of branches. For further details on bifurcation theory, see for instance [4] or specifically in the context of pattern formation [5]. The key mechanism relevant to this paper is depicted in Figure 1. Though the theory is solid, in practice it is typically impossible to calculate a bifurcation diagram in any but the simplest examples. Instead, one uses numerical algorithms termed **continuation** methods [4, 6], whereby discrete points are estimated along the bifurcation diagram one after another. Though the principle may seem simple, effective and reliable continuation methods are challenging to develop especially for large sys-

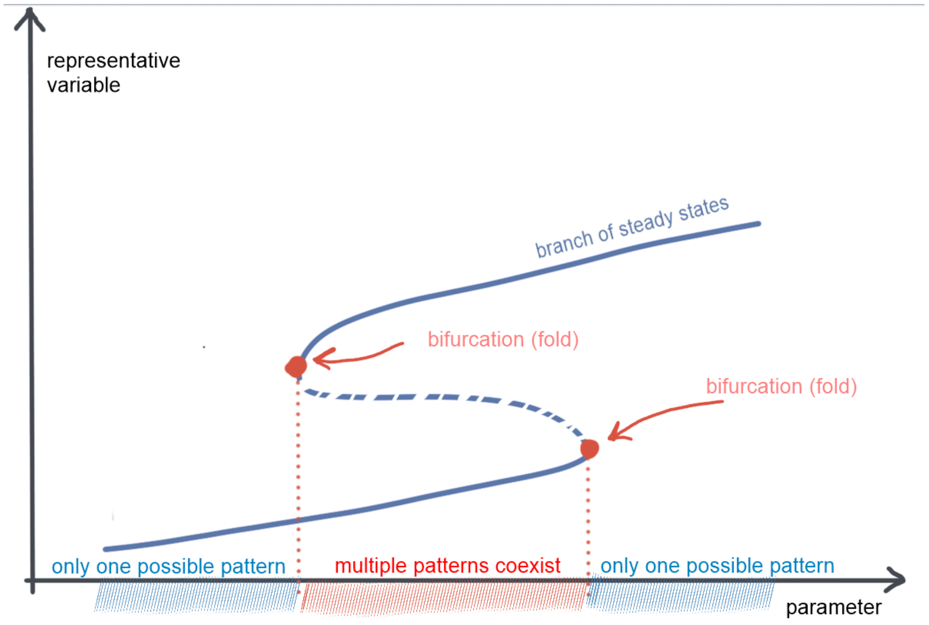


Figure 1: Sketch of a bifurcation diagram including two **fold bifurcations**. In abscissa, an arbitrary parameter. In the intermediary region, there are two stable steady states (solid branches) and an unstable (dashed branch) one in-between. For lower/higher values of the parameter, the system presents a single, stable steady state: the asymptotic dynamics is independent of the initial conditions.

tems of differential equations, as occurs for spatially extended systems (where each spatial location requires at least on differential equation).
