## Supplementary material for "Fluctuations in auxin levels depend upon synchronicity of cell divisions in a one-dimensional model of auxin transport": S2_File_code: continuation.m

```
%code altered from original script by Daniele Avitabile
% Copyright 2016 by Daniele Avitabile
% See https://www.maths.nottingham.ac.uk/personal/pmzda/
%
% If you use this code, please cite
% Daniele Avitabile, "Numerical computation of coherent structures in
% spatially-extended neural networks", Second International Conference on
% Mathematical Neuroscience, Antibes Juan-les-Pins, 2016

%% Clean
clear all, close all, clc;

%% Load folders
addpath('../../../SecantContinuation/0.2/src');
addpath('../ProblemFiles');
addpath('../SolutionDatabase');

%% Spatial coordinates: x direction
bConds = 'periodic';
xcells = 128;
ycells = 1;
nx = xcells*ycells; x = 0 +[0:nx-1]*1;
dimension = 1;

%% Load initial guess and parameters
load('InitialGuess_and_parameters.mat')


%% Get linear operators
[L,N,V] = VolumeAndL_Grid(xcells,ycells,dimension,nx);


%% Define handle to right-hand side and time output function
prob     = @(u,p) Flux2D(u,p,L,N,V);
plotSol  = @(u,p,parent) PlotSolution(x,u(1:nx),p,parent);  
compSpec = @(u,p) ComputeSpectrum2d(u,p,L,V,N);
solMeas  = @(step,u,p) SolutionMeasures(step,u,p,x,bConds);
plotSpec = @(lambda,p,parent) PlotSpectrum(lambda,p,parent);


%% Assign problem
stepPars.iContPar             = 5;
stepPars.pMin                 = -20;
stepPars.pMax                 = 20;
stepPars.s0                   = 0.05; %-0.05; %signed step size
stepPars.sMin                 = 0.0001;
stepPars.sMax                 = 0.1;%0.5; %0.3;%0.05;
stepPars.maxSteps             = 20000;
stepPars.nPrint               = 1;
stepPars.nSaveSol             = 1;
stepPars.finDiffEps           = 1e-7;
stepPars.fsolveOptions        = optimset('Display','iter',...
                                            'TolFun',1e-4,...
                                            'MaxIter',10,...
                                            'Jacobian','on',...
                                            'DerivativeCheck','off');
stepPars.optNonlinIter        = 5;
stepPars.dataFolder           = './Data';
stepPars.PlotSolution         = plotSol;
stepPars.BranchVariables      = solMeas;
stepPars.PlotBranchVariableId = 2;
stepPars.ComputeEigenvalues   = compSpec;
stepPars.PlotSpectrum         = plotSpec;

%% Run
branch = SecantContinuation(prob,u0,p0,stepPars);
```
