## Supplementary material for "Fluctuations in auxin levels depend upon synchronicity of cell divisions in a one-dimensional model of auxin transport": S2_File_code: Cell_division_events.m

```
function [value,isterminal,direction] = Cell_division_events(t,U)
%CELL_DIVISION_EVENTS checks for when a cell is ready to divide so as to
%pause the ode for its resolution
global num_cells

value = double(U(num_cells*2+1:num_cells*3)  >4/3);     % Detect if any cells have sufficient volume to divide
isterminal = ones(num_cells,1);   % States that the integration stops if above condition is met
direction = zeros(length(U(num_cells*2+1:num_cells*3)),1);   % Set to zero, we do not need to differentiate between a cell increaseing or decreasing in volume as the latter is impossible.
end
```
