## Supplementary material for "Fluctuations in auxin levels depend upon synchronicity of cell divisions in a one-dimensional model of auxin transport": S2_File_code: Divide_cells.m

```
function [u,N,L] = Divide_cells(u,N,ie,division_error,t)
%DIVIDE_CELLS handle the cell division event in the growth of the plant
global num_cells
%expand u and insert values for new cell
u = [u(1:num_cells)';u(ie);u(num_cells+1:2*num_cells)';u(ie+num_cells);u(2*num_cells+1:3*num_cells)';u(2*num_cells + ie)/2];
num_cells = num_cells+1;

%update cell ie
%alter cell sizes based on distribution for division if used
if division_error == 1
    difference = random(t,1)/100*u(2*num_cells+ie);
    u(2*num_cells+ie) = u(2*num_cells+ie)/2 + difference;
    u(3*num_cells) = u(3*num_cells) - difference;
else
    u(2*num_cells+ie) = u(2*num_cells+ie)/2 ;
end

    


%handle change of adjacency left and right
N.left(num_cells,:) = [2,ie];
if N.right(ie,2) ~= 0
    N.left(N.right(ie,2),:) = [2,num_cells];
end
N.right(num_cells,:) = N.right(ie,:);
N.right(ie,:) = [2,num_cells];
%update row length
N.row_length(N.position(ie,2)) = N.row_length(N.position(ie,2)) + 1;


%update the N.possition
for i = 1:num_cells-1
    if N.position(i,2) == N.position(ie,2) && N.position(i,1) > N.position(ie,1)
        N.position(i,1) = N.position(i,1) + 1;
    end
end
N.position(num_cells,2) = N.position(ie,2);
N.position(num_cells,1) = N.position(ie,1) + 1;

%update N.self
N.self(num_cells) = 0;

%resort the sorted list

[~,I] = sort(N.position(N.position(:,2) ==N.position(ie,2),1),1);
index = [1:num_cells]';
index = index(N.position(:,2) ==N.position(ie,2),1);
N.position_sorted(N.position(ie,2),1:N.row_length(N.position(ie,2))) = index(I)';

%update N.growing
N.growing(num_cells) = 1;
end
```
