## Supplementary material for "Fluctuations in auxin levels depend upon synchronicity of cell divisions in a one-dimensional model of auxin transport": S2_File_code: Flux2D_diff_and_div.m

```
% Copyright 2016 by Daniele Avitabile
% See https://www.maths.nottingham.ac.uk/personal/pmzda/
%
% If you use this code, please cite
% Daniele Avitabile, "Numerical computation of coherent structures in
% spatially-extended neural networks", Second International Conference on
% Mathematical Neuroascience, Antibes Juan-les-Pins, 2016

function [F,J] = Flux2D_diff_and_div(u,p,N)
%a is a vector storing theconcentration of auxin and the concentration of
%pin protiens in cells
global num_cells


% Rename parameters
n = size(u,1)/3;
rhoIAA = p(1,1);
kIAA = p(2,1);
mIAA = p(3,1);
D = p(4,1);
T = p(5,1);
c1 = p(6,1);
kT = p(7,1);
rhoPIN0 = p(8,1);
rhoPIN = p(9,1);
kPIN = p(10,1);
mPIN = p(11,1);
cell_growth_rate = p(12,1);


%generate cumulative sum lengths
for i = 1:N.num_rows
    cumulative_length(i,1:N.row_length(i)+1) = [0,cumsum(u(N.position_sorted(i,1:N.row_length(i))+2*num_cells)')];
end

%get adjacency matrix
L=sparse(n,n);
%get neighbours
if nargout >1
    for i= 1:num_cells
        if N.self(i) == 1
            L(i,i) = 1;
        end
        N.neighbours(i,1) = 1;
        for j = 2:N.left(i,1)
            N.neighbours(i,1) = N.neighbours(i,1) + 1;
            N.neighbours(i,N.neighbours(i,1)) = N.left(i,j);
            L(i,N.left(i,j)) = 1;
            L(N.left(i,j),i) = 1;
        end
        for j = 2:N.right(i,1)
            N.neighbours(i,1) = N.neighbours(i,1) + 1;
            N.neighbours(i,N.neighbours(i,1)) = N.right(i,j);
            L(i,N.right(i,j)) = 1;
            L(N.right(i,j),i) = 1;
        end
    end
    for i = 1:N.num_rows-1
        %let j be the index for the row above and k be the index for the row
        %below
        j=2;k=2;
        while j<=N.row_length(i)+1 && k<=N.row_length(i+1)+1
            if cumulative_length(i,j) < cumulative_length(i+1,k)
                if cumulative_length(i,j-1) < cumulative_length(i+1,k-1)
                    L(N.position_sorted(i,j-1),N.position_sorted(i+1,k-1)) = cumulative_length(i,j)-cumulative_length(i+1,k-1);
                    L(N.position_sorted(i+1,k-1),N.position_sorted(i,j-1)) = L(N.position_sorted(i,j-1),N.position_sorted(i+1,k-1));
                    N.neighbours(N.position_sorted(i,j-1),1) = N.neighbours(N.position_sorted(i,j-1),1) + 1;
                    N.neighbours(N.position_sorted(i,j-1),N.neighbours(N.position_sorted(i,j-1),1)) = N.position_sorted(i,k-1);
                    N.neighbours(N.position_sorted(i,k-1),1) = N.neighbours(N.position_sorted(i,k-1),1) + 1;
                    N.neighbours(N.position_sorted(i,k-1),N.neighbours(N.position_sorted(i,k-1),1)) = N.position_sorted(i,j-1);
                elseif cumulative_length(i,j-1) >cumulative_length(i+1,k-1)
                    L(N.position_sorted(i,j-1),N.position_sorted(i+1,k-1)) = cumulative_length(i,j)-cumulative_length(i,j-1);
                    L(N.position_sorted(i+1,k-1),N.position_sorted(i,j-1)) = L(N.position_sorted(i,j-1),N.position_sorted(i+1,k-1));
                    N.neighbours(N.position_sorted(i,j-1),1) = N.neighbours(N.position_sorted(i,j-1),1) + 1;
                    N.neighbours(N.position_sorted(i,j-1),N.neighbours(N.position_sorted(i,j-1),1)) = N.position_sorted(i,k-1);
                    N.neighbours(N.position_sorted(i,k-1),1) = N.neighbours(N.position_sorted(i,k-1),1) + 1;
                    N.neighbours(N.position_sorted(i,k-1),N.neighbours(N.position_sorted(i,k-1),1)) = N.position_sorted(i,j-1);
                else
                    L(N.position_sorted(i,j-1),N.position_sorted(i+1,k-1)) = cumulative_length(i,j)-cumulative_length(i,j-1);
                    L(N.position_sorted(i+1,k-1),N.position_sorted(i,j-1)) = L(N.position_sorted(i,j-1),N.position_sorted(i+1,k-1));
                    N.neighbours(N.position_sorted(i,j-1),1) = N.neighbours(N.position_sorted(i,j-1),1) + 1;
                    N.neighbours(N.position_sorted(i,j-1),N.neighbours(N.position_sorted(i,j-1),1)) = N.position_sorted(i,k-1);
                    N.neighbours(N.position_sorted(i,k-1),1) = N.neighbours(N.position_sorted(i,k-1),1) + 1;
                    N.neighbours(N.position_sorted(i,k-1),N.neighbours(N.position_sorted(i,k-1),1)) = N.position_sorted(i,j-1);
                end
                j = j+1;
            elseif cumulative_length(i,j) > cumulative_length(i+1,k)
                if cumulative_length(i,j-1) < cumulative_length(i+1,k-1)
                    L(N.position_sorted(i,j-1),N.position_sorted(i+1,k-1)) = cumulative_length(i+1,k)-cumulative_length(i+1,k-1);
                    L(N.position_sorted(i+1,k-1),N.position_sorted(i,j-1)) = L(N.position_sorted(i,j-1),N.position_sorted(i+1,k-1));
                    N.neighbours(N.position_sorted(i,j-1),1) = N.neighbours(N.position_sorted(i,j-1),1) + 1;
                    N.neighbours(N.position_sorted(i,j-1),N.neighbours(N.position_sorted(i,j-1),1)) = N.position_sorted(i,k-1);
                    N.neighbours(N.position_sorted(i,k-1),1) = N.neighbours(N.position_sorted(i,k-1),1) + 1;
                    N.neighbours(N.position_sorted(i,k-1),N.neighbours(N.position_sorted(i,k-1),1)) = N.position_sorted(i,j-1);
                elseif cumulative_length(i,j-1) >cumulative_length(i+1,k-1)
                    L(N.position_sorted(i,j-1),N.position_sorted(i+1,k-1)) = cumulative_length(i+1,k)-cumulative_length(i,j-1);
                    L(N.position_sorted(i+1,k-1),N.position_sorted(i,j-1)) = L(N.position_sorted(i,j-1),N.position_sorted(i+1,k-1));
                    N.neighbours(N.position_sorted(i,j-1),1) = N.neighbours(N.position_sorted(i,j-1),1) + 1;
                    N.neighbours(N.position_sorted(i,j-1),N.neighbours(N.position_sorted(i,j-1),1)) = N.position_sorted(i,k-1);
                    N.neighbours(N.position_sorted(i,k-1),1) = N.neighbours(N.position_sorted(i,k-1),1) + 1;
                    N.neighbours(N.position_sorted(i,k-1),N.neighbours(N.position_sorted(i,k-1),1)) = N.position_sorted(i,j-1);
                else
                    L(N.position_sorted(i,j-1),N.position_sorted(i+1,k-1)) = cumulative_length(i+1,k)-cumulative_length(i,j-1);
                    L(N.position_sorted(i+1,k-1),N.position_sorted(i,j-1)) = L(N.position_sorted(i,j-1),N.position_sorted(i+1,k-1));
                    N.neighbours(N.position_sorted(i,j-1),1) = N.neighbours(N.position_sorted(i,j-1),1) + 1;
                    N.neighbours(N.position_sorted(i,j-1),N.neighbours(N.position_sorted(i,j-1),1)) = N.position_sorted(i,k-1);
                    N.neighbours(N.position_sorted(i,k-1),1) = N.neighbours(N.position_sorted(i,k-1),1) + 1;
                    N.neighbours(N.position_sorted(i,k-1),N.neighbours(N.position_sorted(i,k-1),1)) = N.position_sorted(i,j-1);
                end
                k = k+1;
            else
                if cumulative_length(i,j-1) < cumulative_length(i+1,k-1)
                    L(N.position_sorted(i,j-1),N.position_sorted(i+1,k-1)) = cumulative_length(i+1,k)-cumulative_length(i+1,k-1);
                    L(N.position_sorted(i+1,k-1),N.position_sorted(i,j-1)) = L(N.position_sorted(i,j-1),N.position_sorted(i+1,k-1));
                    N.neighbours(N.position_sorted(i,j-1),1) = N.neighbours(N.position_sorted(i,j-1),1) + 1;
                    N.neighbours(N.position_sorted(i,j-1),N.neighbours(N.position_sorted(i,j-1),1)) = N.position_sorted(i,k-1);
                    N.neighbours(N.position_sorted(i,k-1),1) = N.neighbours(N.position_sorted(i,k-1),1) + 1;
                    N.neighbours(N.position_sorted(i,k-1),N.neighbours(N.position_sorted(i,k-1),1)) = N.position_sorted(i,j-1);
                elseif cumulative_length(i,j-1) >cumulative_length(i+1,k-1)
                    L(N.position_sorted(i,j-1),N.position_sorted(i+1,k-1)) = cumulative_length(i+1,k)-cumulative_length(i,j-1);
                    L(N.position_sorted(i+1,k-1),N.position_sorted(i,j-1)) = L(N.position_sorted(i,j-1),N.position_sorted(i+1,k-1));
                    N.neighbours(N.position_sorted(i,j-1),1) = N.neighbours(N.position_sorted(i,j-1),1) + 1;
                    N.neighbours(N.position_sorted(i,j-1),N.neighbours(N.position_sorted(i,j-1),1)) = N.position_sorted(i,k-1);
                    N.neighbours(N.position_sorted(i,k-1),1) = N.neighbours(N.position_sorted(i,k-1),1) + 1;
                    N.neighbours(N.position_sorted(i,k-1),N.neighbours(N.position_sorted(i,k-1),1)) = N.position_sorted(i,j-1);
                else
                    L(N.position_sorted(i,j-1),N.position_sorted(i+1,k-1)) = cumulative_length(i+1,k)-cumulative_length(i,j-1);
                    L(N.position_sorted(i+1,k-1),N.position_sorted(i,j-1)) = L(N.position_sorted(i,j-1),N.position_sorted(i+1,k-1));
                    N.neighbours(N.position_sorted(i,j-1),1) = N.neighbours(N.position_sorted(i,j-1),1) + 1;
                    N.neighbours(N.position_sorted(i,j-1),N.neighbours(N.position_sorted(i,j-1),1)) = N.position_sorted(i,k-1);
                    N.neighbours(N.position_sorted(i,k-1),1) = N.neighbours(N.position_sorted(i,k-1),1) + 1;
                    N.neighbours(N.position_sorted(i,k-1),N.neighbours(N.position_sorted(i,k-1),1)) = N.position_sorted(i,j-1);
                end
                k = k+1;
                j = j+1;
            end
        end
    end
else
    for i= 1:num_cells
        if N.self(i) == 1
            L(i,i) = 1;
        end
        N.Neighbours(i,1) = 1;
        for j = 2:N.left(i,1)
            L(i,N.left(i,j)) = 1;
            L(N.left(i,j),i) = 1;
        end
        try
        for j = 2:N.right(i,1)
            L(i,N.right(i,j)) = 1;
            L(N.right(i,j),i) = 1;
        end
        catch
            i
            j
        end
    end
    for i = 1:N.num_rows-1
        %let j be the index for the row above and k be the index for the row
        %below
        j=2;k=2;
        while j<=N.row_length(i)+1 && k<=N.row_length(i+1)+1
            if cumulative_length(i,j) < cumulative_length(i+1,k)
                if cumulative_length(i,j-1) < cumulative_length(i+1,k-1)
                    L(N.position_sorted(i,j-1),N.position_sorted(i+1,k-1)) = cumulative_length(i,j)-cumulative_length(i+1,k-1);
                    L(N.position_sorted(i+1,k-1),N.position_sorted(i,j-1)) = L(N.position_sorted(i,j-1),N.position_sorted(i+1,k-1));
                    
                elseif cumulative_length(i,j-1) >cumulative_length(i+1,k-1)
                    L(N.position_sorted(i,j-1),N.position_sorted(i+1,k-1)) = cumulative_length(i,j)-cumulative_length(i,j-1);
                    L(N.position_sorted(i+1,k-1),N.position_sorted(i,j-1)) = L(N.position_sorted(i,j-1),N.position_sorted(i+1,k-1));
                    
                else
                    L(N.position_sorted(i,j-1),N.position_sorted(i+1,k-1)) = cumulative_length(i,j)-cumulative_length(i,j-1);
                    L(N.position_sorted(i+1,k-1),N.position_sorted(i,j-1)) = L(N.position_sorted(i,j-1),N.position_sorted(i+1,k-1));
                    
                end
                j = j+1;
            elseif cumulative_length(i,j) > cumulative_length(i+1,k)
                if cumulative_length(i,j-1) < cumulative_length(i+1,k-1)
                    L(N.position_sorted(i,j-1),N.position_sorted(i+1,k-1)) = cumulative_length(i+1,k)-cumulative_length(i+1,k-1);
                    L(N.position_sorted(i+1,k-1),N.position_sorted(i,j-1)) = L(N.position_sorted(i,j-1),N.position_sorted(i+1,k-1));
                    
                elseif cumulative_length(i,j-1) >cumulative_length(i+1,k-1)
                    L(N.position_sorted(i,j-1),N.position_sorted(i+1,k-1)) = cumulative_length(i+1,k)-cumulative_length(i,j-1);
                    L(N.position_sorted(i+1,k-1),N.position_sorted(i,j-1)) = L(N.position_sorted(i,j-1),N.position_sorted(i+1,k-1));
                    
                else
                    L(N.position_sorted(i,j-1),N.position_sorted(i+1,k-1)) = cumulative_length(i+1,k)-cumulative_length(i,j-1);
                    L(N.position_sorted(i+1,k-1),N.position_sorted(i,j-1)) = L(N.position_sorted(i,j-1),N.position_sorted(i+1,k-1));
                    
                end
                k = k+1;
            else
                if cumulative_length(i,j-1) < cumulative_length(i+1,k-1)
                    L(N.position_sorted(i,j-1),N.position_sorted(i+1,k-1)) = cumulative_length(i+1,k)-cumulative_length(i+1,k-1);
                    L(N.position_sorted(i+1,k-1),N.position_sorted(i,j-1)) = L(N.position_sorted(i,j-1),N.position_sorted(i+1,k-1));
                    
                elseif cumulative_length(i,j-1) >cumulative_length(i+1,k-1)
                    L(N.position_sorted(i,j-1),N.position_sorted(i+1,k-1)) = cumulative_length(i+1,k)-cumulative_length(i,j-1);
                    L(N.position_sorted(i+1,k-1),N.position_sorted(i,j-1)) = L(N.position_sorted(i,j-1),N.position_sorted(i+1,k-1));
                    
                else
                    L(N.position_sorted(i,j-1),N.position_sorted(i+1,k-1)) = cumulative_length(i+1,k)-cumulative_length(i,j-1);
                    L(N.position_sorted(i+1,k-1),N.position_sorted(i,j-1)) = L(N.position_sorted(i,j-1),N.position_sorted(i+1,k-1));
                    
                end
                k = k+1;
                j = j+1;
            end
        end
    end
end
%pin protein matrix

P =    L.* (((u(n+1:2*n)  ./  (L*exp(c1*u(1:n)))))   *  transpose(((exp(c1*u(1:n))))));
dPdp = L.* (((1           ./  (L*exp(c1*u(1:n)))))   *  transpose(((exp(c1*u(1:n))))));
dPda = L.* (((u(n+1:2*n)  ./  (L*exp(c1*u(1:n))).^2)) *  transpose(((exp(c1*u(1:n))))));
V = u(2*n+1:3*n);

F = zeros(2*n,1);
% Right-hand side
F(1:n,1) = rhoIAA./(1+ kIAA.* u(1:n)) - mIAA.*u(1:n) - cell_growth_rate*u(1:n)./V(1:n) ...
    + (D./V).*(L*u(1:n) - sum(L,2).*u(1:n)) ...
    + (T./V).*( transpose(P)*(u(1:n).^2).*(1./(1+kT.*(u(1:n).^2))) ...
    - P*(1./(1 + kT*u(1:n).^2)) .* (u(1:n).^2)) ;
F(n+1:2*n) = (rhoPIN0 + rhoPIN.* u(1:n))./(1+ kPIN.* u(n+1:2*n)) - mPIN.*u(n+1:2*n) - cell_growth_rate*u(n+1:2*n)./V(1:n);
% F(1:n/2,1) = F(1:n/2,1) + epsilon;
% Jacobian
F(2*n+1:3*n) = cell_growth_rate*N.growing;


if nargout > 1
    J = zeros(2*n);
    
    for i = 1:n
        
        %first term
        J(i,i) = - (kIAA*rhoIAA)/(1+kIAA*u(i))^2 - mIAA;
        
        %second term
        for j = 3:N(i,1)
            J(i,N(i,j)) = J(i,N(i,j)) + D/V(i)*L(i,N(i,j));
            J(i,i)      = J(i,i)      - D/V(i)*L(i,N(i,j));
        end
        
        %third term
        for j = 3:N(i,1)
            
            %third term part 1
            for k = 3:N(N(i,j),1)
                J(i,N(N(i,j),k)) = J(i,N(N(i,j),k)) - (T/V(i))*c1*L(N(i,j),N(N(i,j),k))*exp(c1*u(N(N(i,j),k))) * dPda(N(i,j),i) * (u(N(i,j))^2/(1 + kT*u(i)^2));
            end
            J(i,i) = J (i,i) + (T/V(i))*c1 * P(N(i,j),i) *(u(N(i,j))^2/(1 + kT*u(i)^2)) - (T/V(i))*P(N(i,j),i)*(2*kT*u(i)*u(N(i,j))^2/(1 + kT*u(i)^2)^2);
            if N(N(i,j),2) == 1
                J(i,N(i,j)) = J(i,N(i,j)) - (T/V(i))*c1*L(N(i,j),N(i,j))*exp(c1*u(N(i,j))) * dPda(N(i,j),i) * (u(N(i,j))^2/(1 + kT*u(i)^2)) + (T/V(i))*P(N(i,j),i)*2*(u(N(i,j))/(1 + kT*u(i)^2));
            else
                J(i,N(i,j)) = J(i,N(i,j)) + (T/V(i))*P(N(i,j),i)*2*(u(N(i,j))/(1 + kT*u(i)^2));
            end
            
            %third term part 2
            for k = 3:j-1
                J(i,N(i,k)) = J(i,N(i,k)) + (T/V(i))*c1*L(i,N(i,k))*exp(c1*u(N(i,k))) * dPda(i,N(i,j)) * (u(i)^2/(1 + kT*u(N(i,j))^2));
            end
            J(i,N(i,j)) = J(i,N(i,j)) - (T/V(i))*(c1*P(i,N(i,j)) -  c1*L(i,N(i,j))*exp(c1*u(N(i,j))) * dPda(i,N(i,j))) *(u(i)^2/(1 + kT*u(N(i,j))^2))  + (T/V(i))*P(i, N(i,j))* 2*kT*u(N(i,j))*u(i)^2 / (1 + kT*u(N(i,j))^2)^2  ;
            for k = j+1:N(i,1)
                J(i,N(i,k)) = J(i,N(i,k)) + (T/V(i))*c1*L(i,N(i,k))*exp(c1*u(N(i,k))) * dPda(i,N(i,j)) * (u(i)^2/(1 + kT*u(N(i,j))^2));
            end
            if N(i,2) == 1
                J(i,i) = J(i,i)  - (T/V(i))*P(i,N(i,j))*(2*u(i)/(1 + kT*u(N(i,j))^2))+ (T/V(i))*c1*L(i,i)*exp(c1*u(i)) * dPda(i,N(i,j)) * (u(i)^2/(1 + kT*u(N(i,j))^2))    ;
            else
                J(i,i) = J(i,i)  - (T/V(i))*P(i,N(i,j))*(2*u(i)/(1 + kT*u(N(i,j))^2));%+ (T/V(i))*c1*L(i,N(i,k))*exp(c1*u(N(i,k))) * dPda(i,N(i,j)) * (u(i)^2/(1 + kT*u(N(i,j))^2))
            end
            
            
            %third term with respect to p
            
            J(i,N(i,j) + n) =            (T/V(i))*dPdp(N(i,j),i) * (u(N(i,j))^2/(1 + kT*u(i)^2));
            J(i,i+n)        = J(i,i+n) - (T/V(i))*dPdp(i,N(i,j)) * (u(i)^2/(1 + kT*u(N(i,j))^2));
            
        end
    end
    
    for i = n+1:2*n
        J(i,i-n) = rhoPIN/(1 + kPIN*u(i));
        J(i,i) = -( kPIN*(rhoPIN0 + rhoPIN*u(i-n)))/((1 + kPIN*u(i))^2) - mPIN;
    end
end

end
```
