## Supplementary material for "Fluctuations in auxin levels depend upon synchronicity of cell divisions in a one-dimensional model of auxin transport": S2_File_code: PlotHistory_fixedtip.m

```
function [] = PlotHistory_fixedtip(U,N)
global num_cells
num_cells = size(U,2)/3;
root_length = floor(10*sum(U(end,(2*num_cells+1):(3*num_cells)))) +1;
number_of_time_steps = size(U,1);
history_plot = nan(number_of_time_steps,root_length);
for i = 1:number_of_time_steps
        cumsum_rowlength = floor(10*[0,cumsum(U(i,(N.position_sorted+2*num_cells)))])+1;
        %adjust to put root tip at end of plot
        cumsum_rowlength = cumsum_rowlength + (root_length -cumsum_rowlength(end))+1;
        
        for k = 1:length(N.position_sorted)
            try
            history_plot(i,cumsum_rowlength(k):cumsum_rowlength(k+1)) = U(i,N.position_sorted(k));
            catch
                poop = 2
            end
        end
end
fixedtipauxin = nan(number_of_time_steps,num_cells);
for i = 1:number_of_time_steps
    auxinrow = U(i,N.position_sorted);
    nonzeroauxinrow = auxinrow(auxinrow~=0);
    
    fixedtipauxin(i,(num_cells-length(nonzeroauxinrow)+1):num_cells) = nonzeroauxinrow;
end


figure
auxinhistory = image(history_plot,'CDataMapping','scaled')
set(auxinhistory,'AlphaData',~isnan(history_plot))
caxis([min(U(end,1:num_cells)),max(U(end,1:num_cells))])
caxis(([min(U(end,1:num_cells)),max(U(end,1:num_cells))] + mean(U(end,1:num_cells)))/2)
caxis([1,3])
end
```
