## Supplementary material for "Fluctuations in auxin levels depend upon synchronicity of cell divisions in a one-dimensional model of auxin transport": S2_File_code: TimeOutput.m

```
% Copyright 2016 by Daniele Avitabile
% See https://www.maths.nottingham.ac.uk/personal/pmzda/
%
% If you use this code, please cite
% Daniele Avitabile, "Numerical computation of coherent structures in
% spatially-extended neural networks", Second International Conference on
% Mathematical Neuroscience, Antibes Juan-les-Pins, 2016

function status = TimeOutput(t,u,flag)

  if isempty(flag)
    disp(['t = ' num2str(max(t))]);
  end
  status = 0;

end
```
