## Supplementary material for "Fluctuations in auxin levels depend upon synchronicity of cell divisions in a one-dimensional model of auxin transport": S2_File_code: VolumeAndL_for_div.m

```
function [N] = VolumeAndL_for_div(xcells)
%VOLUMEANDL Summary of this function goes here

% N.up = [#entries,neighours above in left to right order]
% n.left ...
% n.right ...
% n.down ...
% n.self = [1 if references self
% n.row = # of the row for graphing

%dimesions for up and down and row are given due to rest of of code
%allowing expansion to 2d cell geometries


    N.up(1:xcells,1) = ones(xcells,1);
    N.left(1:xcells,1) = [1; 2*ones(xcells-1,1)]; N.left(1:xcells,2) = [0; (1:xcells-1)'];
    N.right(1:xcells,1) = [2*ones(xcells-1,1); 1]; 
    N.right(1:xcells,2) = [(2:xcells)';0];
    N.down(1:xcells,1) = ones(xcells,1);
    N.self = [1;zeros(xcells-1,1)];
    N.position = [(1:xcells)',ones(xcells,1);inf*ones(xcells-xcells,1),inf*ones(xcells-xcells,1)];
                   
    N.row_length = xcells;
    N.position_sorted = [1:xcells];
    N.num_rows = 1;
    N.row_length = xcells;
    N.growing = ones(xcells,1);

end
```
