## Supplementary material for "Fluctuations in auxin levels depend upon synchronicity of cell divisions in a one-dimensional model of auxin transport": S2_File_code: timeStep.m

```
%code heavily altered from original script by Daniele Avitabile
% Copyright 2016 by Daniele Avitabile
% See https://www.maths.nottingham.ac.uk/personal/pmzda/
%
% If you use this code, please cite
% Daniele Avitabile, "Numerical computation of coherent structures in
% spatially-extended neural networks", Second International Conference on
% Mathematical Neuroscience, Antibes Juan-les-Pins, 2016

% Cleaning 
clear all, close all, clc;

% Load problem files and solution
addpath('../ProblemFiles');

% Spatial coordinates: x direction
xcells = 2;
ycells = 1;
%nx = 100; Lx = 50; hx = 2*Lx/nx; x = 0 +[0:nx-1]*1;
global num_cells
num_cells = ycells*xcells;
%number of cells at the tip that are able to divide
growth_area = 8;
 

 tend = 2000;
 tincrement = 1;
% Get linear operators
[N] = VolumeAndL_for_div(xcells);

%Set parameters
  p0(1,1) = 0.85; %rhoIAA
  p0(2,1) = 1; %kIAAShanties: All wooden ships leak somewhat. There was a special hold (cargo area) in the ships wh
  p0(3,1) = 0.1; %mIAA
  p0(4,1) = 1;%0.00003;%1.00; %D
  p0(5,1) = 2.1;%1.9; %T
  p0(6,1) = 1.099; %c10
  p0(7,1) = 1; %kT
  p0(8,1) = 0; %rhoPIN0
  p0(9,1) = 1; %rhoPIN
  p0(10,1) = 1; %kPIN
  p0(11,1) = .1;%.1;%0.1; %mPIN
  p0(12,1) =10/3000; %1.0;%theta
  p0(13,1) = 1; %rho
  p0(14,1) = 0; %alpha
  p0(15,1) = 1; %LbyW

  
%set up the distribution used for stochastic cell division sizes 
%for a normal ditribution use :
pd = makedist('Normal','mu',0,'sigma',9.3);
truncatednormal = truncate(pd,-30,30); 
Distribution = truncatednormal;

% %for a uniform distribution use:
% Uniformdist = makedist('Uniform','lower',-10,'upper',10);
% Distribution = Uniformdist;

% set to 1 if using a distribution or 0 if cells divide exactly in two
division_error = 1;
%initialise vector recording times for cell division events
celldivisiontime = [];

%initialise the auxin, pins and size of cells
 if num_cells > 16
    u0 = [20/15*ones(num_cells - 16,1);10/15*ones(16,1)]; u0 = [u0;u0;u0;];%20/15*ones(num_cells*3,1);
 else
    u0 = 10/15*ones(num_cells*3,1);
 end
 
 
% % Define handle to right-hand side and time output function
prob = @(t,u) Flux2D_diff_and_div(u,p0,N);

% set ode solver to inform user of progress of simulation
timeOut = @(t,u,flag) TimeOutput(t,u,flag);
Ns = N;
%% Time step
complete = 0;
    tout = [];
    Uout = [];
    teout = [];
    Ueout = [];
    ieout = [];
   tSpan = ([0:tincrement:tend+1]); opts = odeset('OutputFcn',timeOut,'Events',@Cell_division_events);%,'RelTol',1e-20, 'AbsTol', 1e-20);

    [t,U,te,Ue,ie] = ode23s(prob,tSpan,u0,opts);    %%%%%%%%%%%%%%%%%%%% ODE
    %check if tend is reached, if not iterate after resolving cell
    %divisions
    if t(end) > tend
        complete = 1;
    end
    
        %accumulate output
    nt = length(t);
    tout = t(2:nt);
    Uout =  U(2:nt,:);
    teout = te;          % Events at tstart are never reported.
    Ueout = Ue;
    ieout = ie;
    %%
%check if the end time has been reached
while complete == 0
    
    

    
    
    %handle the cell division,
    %calculate how many cells are dividing
    num_newcells = 0;
    num_events = length(ie);
    %get cell information at this time
    u0 = U(end,:)';
    
    
    dividingcells = [];
    for i = 1:num_events
        %check if cell that reached division size is within the growth area
        if N.position(ie(i),1) > N.row_length(N.position(ie(i),2)) -growth_area
            %if in growth area add to list of cells to divide
            dividingcells = [dividingcells;i];
        else
            % if not within growth area stop the cell growing further
            N.growing(ie(i)) = 0;
        end
    end
    
    %divide each of the cells and note the time in teh celldivisiontime
    %vector
    for i = 1:length(dividingcells)
        [u0,N] = Divide_cells(u0',N,ie(dividingcells(i)),division_error,Distribution);
        celldivisiontime = [celldivisiontime,t(end)];
        num_newcells = num_newcells+1;
    end

%Save a snapshot of the cell structure
Ns(length(u0)/3).up = N.up;
Ns(length(u0)/3).left = N.left;
Ns(length(u0)/3).right = N.right;
Ns(length(u0)/3).down = N.down;
Ns(length(u0)/3).position = N.position;
Ns(length(u0)/3).row_length = N.row_length;
Ns(length(u0)/3).position_sorted = N.position_sorted;
Ns(length(u0)/3).num_rows = N.num_rows;
Ns(length(u0)/3).growing = N.growing;


    %update callable function with new adjacency information
    prob = @(t,u) Flux2D_diff_and_div(u,p0,N);
    %update time in which next timestepping occures
    tSpan = ([t(end):tincrement:tend+1]);
    [t,U,te,Ue,ie] = ode23s(prob,tSpan,u0,opts); %%%%%%%%%%%%%% ODE
    if t(end) > tend
        %if end time has been reached then times setpping is complete and
        %the loop can be exited
        complete = 1;
    end
    try
         %accumulate output
        nt = length(t);
        tout = [tout; t(2:nt)];
        Uout = [Uout(:,1:num_cells-num_newcells),zeros(size(Uout,1),num_newcells),Uout(:,num_cells-num_newcells+1:2*num_cells-2*num_newcells),zeros(size(Uout,1),num_newcells),Uout(:,2*num_cells-2*num_newcells+1:3*num_cells-3*num_newcells),zeros(size(Uout,1),num_newcells); U(2:nt,:)];
        teout = [teout; te];          % Events at tstart are never reported.
        Ueout = [Ueout(:,1:num_cells-num_newcells),zeros(size(Ueout,1),num_newcells),Ueout(:,num_cells-num_newcells+1:2*num_cells-2*num_newcells),zeros(size(Ueout,1),num_newcells),Ueout(:,2*num_cells-2*num_newcells+1:3*num_cells-3*num_newcells),zeros(size(Ueout,1),num_newcells); Ue];
        %Ueout = [Ueout(:,1:num_cells-1),zeros(size(Ueout,1),1),Ueout(:,num_cells:2*num_cells-2),zeros(size(Ueout,1),1),Ueout(:,2*num_cells-1:3*num_cells-3),zeros(size(Ueout,1),1); Ue];
        ieout = [ieout; ie];
    catch
        %accumulate output
        nt = length(t);
        tout = [tout; t(2:nt)];
        Uout = [Uout; U(2:nt,:)];
    end
end
%%
% Plot history
%set this value to 1 if the tip is to be fixed in space and 0 otherwise.
fix_root_tip = 1;

if fix_root_tip == 1
    PlotHistory_fixedtip(Uout,N)
else
    PlotHistory(Uout,N)
end
```
